# Dendrimer-like supramolecular assembly of proteins with a tunable size and valency through stepwise iterative growth

**DOI:** 10.1101/2021.06.23.449676

**Authors:** Jin-Ho Bae, Hong-Sik Kim, Gijeong Kim, Ji-Joon Song, Hak-Sung Kim

## Abstract

The assembly of proteins in a programmable manner provides insight into the underlying mechanisms of protein self-assembly in nature as well as the creation of novel functional nanomaterials for practical applications. Despite many advances, however, a rational protein assembly with an easy scalability in terms of size and valency remains a challenging task. Here, we present a simple bottom-up approach to the supramolecular protein assembly with a tunable size and valency in a programmable manner. The dendrimer-like protein assembly, called a “prodrimer,” was constructed using a total of three monomeric proteins: a core and two building-block proteins. The prodrimer generations were grown by a stepwise and alternate addition of a building block using two pairs of orthogonal protein-peptide interactions, leading to a higher-generation prodrimer with a mega-dalton size and multi-valency. The valency of the prodrimers at the periphery was tunable with the generation, enabling a single-step functionalization. A second-generation prodrimer functionalized with a target-specific protein binder showed a three-order of magnitude increase in binding affinity compared to a monomeric counterpart due to the avidity. The prodrimers functionalized with a targeting moiety and a cytotoxic protein cargo exhibited a highly enhanced cellular cytotoxicity, exemplifying their utility as a protein delivery platform. The present approach can be effectively used in the creation of protein architectures with new functions for biotechnological and medical applications.

## 1. Introduction

Proteins are the most diverse among functional biomolecules and are building blocks of diverse cellular machineries that perform fundamental cellular tasks. In nature, protein assemblies with higher-order architectures confer remarkable multitudes of new functions that would be impossible when in monomeric forms in terms of functional modulation, allosteric regulation, high structural complexity, and stability ^[1, 2]^. In this regard, a protein assembly has attracted considerable attention as a versatile platform for the creation of highly sophisticated functional nanostructures with novel properties for practical applications in wide-ranging fields, including vaccines, drug delivery, biosensors, diagnosis, and therapy of diseases ^[3–7]^.

A programmable assembly of proteins can provide crucial insight into understanding the underlying mechanisms for protein self-assembly in nature as well as generating unprecedented architectures in a predictable way. The programmability of protein assembly is especially critical for medical and biotechnological purposes where the size and valency of the assembly are the key factors ^[8–10]^. Although proteins have been considered promising building blocks for an assembly owing to their unique structural and biophysical features, the extent of the assembly is highly dependent on the characteristics of the monomeric protein, making precise control of the size and valency difficult. Many strategies for assembling proteins into higher-order structures have been reported, including the use of well-defined coiled-coil and helical bundle interactions, the use of peptide-ligand interactions, the formation of disulfide bonds, chemical cross-links, metal-directed interactions, the use of non-biological templates, and the genetic fusion of self-associating protein domains or fragments ^[5, 11–20]^. The computational design of a protein–protein interface has also recently been applied to the generation of self-assembling proteins with defined structures ^[21–23]^. Despite numerous advances, however, the development of a programmable protein assembly with easy scalability in terms of size and valency remains a challenging task owing to the structural diversity, conformational heterogeneity, and high molecular weight of proteins compared to other biomolecules.

Herein, we describe a bottom-up approach to the supramolecular assembly of proteins with a tunable size and valency in a programmable manner by orthogonal protein-ligand interactions. Coined a “prodrimer,” a dendrimer-like protein assembly is constructed through a stepwise iterative approach using a total of three monomeric proteins (**Scheme 1**): a core (pG_0_) and two building-block proteins (B_1_ and B_2_). With two different isopeptide bond-forming protein and peptide pairs, namely, SpyCatcher/SpyTag and SnoopCatcher/SnoopTag, the generations of prodrimers were grown by an iterative and alternate addition of a building block, allowing a higher-generation prodrimer with a near mega-dalton size and multi-valency. The prodrimers conferred a tunable valency with the generation, leading to a single-step easy functionalization, and their functionalization with a target-specific protein binder demonstrated a typical feature of multi-valency in the avidity and cooperativity. Furthermore, the prodrimers functionalized with a cytotoxic protein cargo exemplified their utility as a protein delivery platform by showing a highly enhanced cellular cytotoxicity compared to a monomeric equivalent. Details are reported herein.

## 2. Results

### 2.1. Strategy for the assembly of dendrimer-like protein architectures

The general procedure for the dendrimer-like protein assembly, coined a “prodrimer,” using a total of three monomeric proteins (a core and two building-block proteins) is depicted in Scheme 1. The core protein (pG_0_, a zeroth-generation prodrimer) consists of a genetically fused dual-tandem SnoopCatcher using a flexible GS linker, whereas the two building blocks (B_1_ and B_2_) comprise dual-tandem repeats of SpyCatcher or SnoopCatcher linked with SnoopTag or SpyTag, respectively (Scheme 1a). Such proteins are a set of orthogonal pairs that spontaneously form a covalent isopeptide bond between a Catcher and a Tag ^[24–27]^. Because each building block is composed of a dissimilar pair of a Catcher and a Tag, no reconstitution or self-cyclization will occur by the single protein alone.

As shown in **Scheme 1b**, the growth of a prodrimer starts with the addition of the building block B_1_ containing an inter SnoopTag to the core protein pG_0_. This results in a first-generation prodrimer, pG_1_, with four SpyCatcher proteins at the periphery through the interaction between SnoopCatcher and SnoopTag. The generation of the prodrimer is further grown by adding the building block B_2_ with an inter SpyTag to pG_1_, yielding a second-generation prodrimer pG_2_ which has a total of eight exposed SnoopCatcher proteins. As the periphery of pG_2_ is returned to SnoopCatcher, the same as pG_0_, the building block B_1_ is added again for the construction of a third-generation prodrimer pG_3_. A fourth-generation prodrimer pG_4_ is produced by adding the building block B_2_ to pG_3_, and consequently pG_4_ has 32 SpyCatcher proteins at the periphery. Similarly, the two building blocks, B_1_ and B_2_, are repetitively added alternatively for the production of a next-generation prodrimer. From only two small building block proteins, prodrimers can be grown to a supramolecular scale in a stepwise manner. Starting from pG_0_ of 27.3 kDa, the prodrimer is sequentially grown to pG_1_, pG_2_, pG_3_, to pG_4_ with a theoretical molecular mass of 92.5, 212.0, 472.6, and 950.5 kDa, respectively. The valency of the prodrimers at the periphery increases by a factor of two after each generation, and consequently pG_4_ has a valency of 32 starting from pG_0_ with a valency of two.

### 2.2. Construction and characterization of prodrimers

Prodrimers are constructed through a bottom-up approach, and it is crucial for each generation to be highly homogeneous. A single malformation in an earlier generation will cause a much greater heterogeneity in the subsequent growth. We thus purified the core protein (pG_0_) and two building blocks (B_1_ and B_2_) to homogeneity by size exclusion chromatography and used them for the construction of prodrimers (Figure S1, Supporting Information).The core and two building block proteins were shown to exhibit a distinct single band at near the expected molecular mass on SDS-PAGE (3%–12% gradient). Following the Scheme 1b, a previous generation prodrimer was purified and used for the construction of a next generation prodrimer using the two building blocks. The iterative formation of different prodrimer generations was traced through size exclusion chromatography (SEC) (Figure S2, Supporting Information). The previous generation prodrimers were shown to be almost completely grown to the next prodrimer generation by alternate addition of a building block. This can be attributed to an efficient step-wise assembly of proteins through specific interactions between orthogonal pairs of SpyCatcher/SpyTag and SnoopCatcher/SnoopTag.

We investigated the biophysical properties of different prodrimer generations after eluted fractions from SEC were collected (**Figure 1**). The prodrimers were observed to exhibit a distinct single band at near the expected molecular mass on SDS-PAGE (**Figure 1a**). The subtle variation in the molecular mass of each generation prodrimers on SDS-PAGE seems to be due to their branched structures even after boiling. In particular, the band of pG_4_ appeared to be slightly smeared mainly owing to the fluid-like low gel percentage at the top. Next, different generations of prodrimers were analyzed based on size-exclusion chromatography, and clear elution peaks were observed without aggregation patterns (**Figure 1b**). DLS analysis also displayed narrow size distributions of the prodrimers, and their average hydrodynamic radii were estimated to range from 5.3 to 20.6 nm (**Figure 1c, 1e**). The absolute molecular masses of the prodrimers were determined using multi-angle light scattering (MALS) **(Figure 1d)**, which were well matched with the expected ones (**Figure 1e)**. Based on the results, it is likely that the present approach enabled supramolecular protein assemblies, i.e., prodrimers, with a well-defined size and valency in a programmable manner.

**Figure 1.**
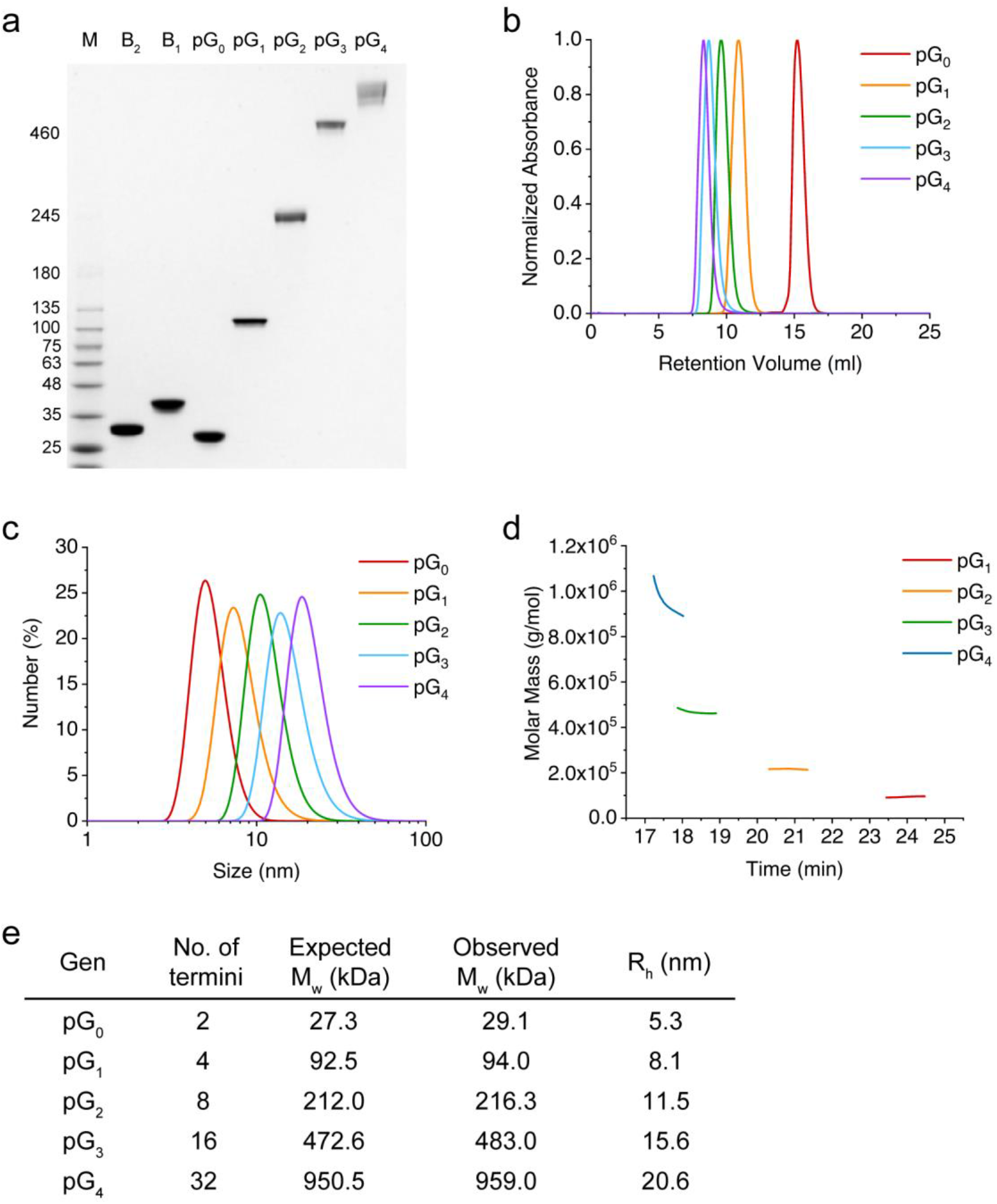
Biophysical characterization of different prodrimer generations. (**a**) SDS-PAGE (3%–12% gradient) of the purified prodrimers from generation 1 (pG_1_) to generation 4 (pG_4_), core protein (pG_0_), and two building block proteins (B_1_ and B_2_). M indicates size markers. (**b**) Size exclusion chromatography (SEC) of different prodrimer generations. The peaks represent the normalized absorbance of different prodrimer generations at 280 nm. (**c**) Dynamic light scattering (DLS) analysis of the prodrimers. (**d**) Absolute molecular mass of the prodrimers by multi-angle light scattering (MALS). The prodrimers eluted from SEC were analyzed. (**e**) Summary of the absolute molecular mass of the prodrimers determined by MALS in (d). The hydrodynamic radii of the prodrimers were determined from the DLS analysis shown in (c).

### 2.3. Functionalization of prodrimers

In order for the prodrimers to be used for biotechnological and medical purposes, their functionalization with biomolecules is essential. With either SpyCatcher or SnoopCatcher at the periphery depending on the generation, we reasoned that the prodrimers could be easily functionalized with diverse biomolecules through interactions with SpyTag or SnoopTag, respectively. We first intended to functionalize the prodrimers for tumor-cell targeting, and employed an EGFR-specific repebody which was previously developed from a non-antibody protein scaffold ^[28, 29]^. For a simple and general functionalization process, a conjugation module comprising of a tandem of SpyTag and SnoopTag was genetically fused to the C-terminal of the repebody using a flexible linker for optimal exposure to yield T_mono_ (indicating a conjugation module-fused targeting moiety) (Figure 2a). As proof-of-concept,we functionalized pG_0_, which has a valency of 2, with T_mono_, and the resulting prodrimer was designated as pG_0_T (standing for a zeroth-generation prodrimer functionalized with T_mono_). Similarly, pG_1_ and pG_2_ were functionalized, yielding pG_1_T and pG_2_T, respectively. Functionalization of different prodrimer generations with a conjugation module-fused EGFR-specific repebody was traced through SEC (Figure S3, Supporting Information).The different generations of prodrimers were observed to be efficiently functionalized, which seems to stem from the fact the conjugation module shares the same chemistry. Analysis of the functionalized prodrimers on SDS-PAGE showed distinct single bands (Figure 2b). From the monomeric T_mono_ with a molecular mass of 32.8 kDa, the molecular mass of pG_2_T increased to approximately 474.6 kDa (Figure 2c). The functionalized prodrimers were eluted with high homogeneity though a size exclusion chromatography, exhibiting a narrow size distribution when analyzed using DLS (Figure S4a, b, Supporting Information). The absolute molecular masses determined through MALS were shown to coincide well with the expected ones (Figure 2c; Figure S4c, Supporting Information). These results demonstrate that the prodrimers can be functionalized using the conjugation module in a highly specific and efficient manner.

**Figure 2.**
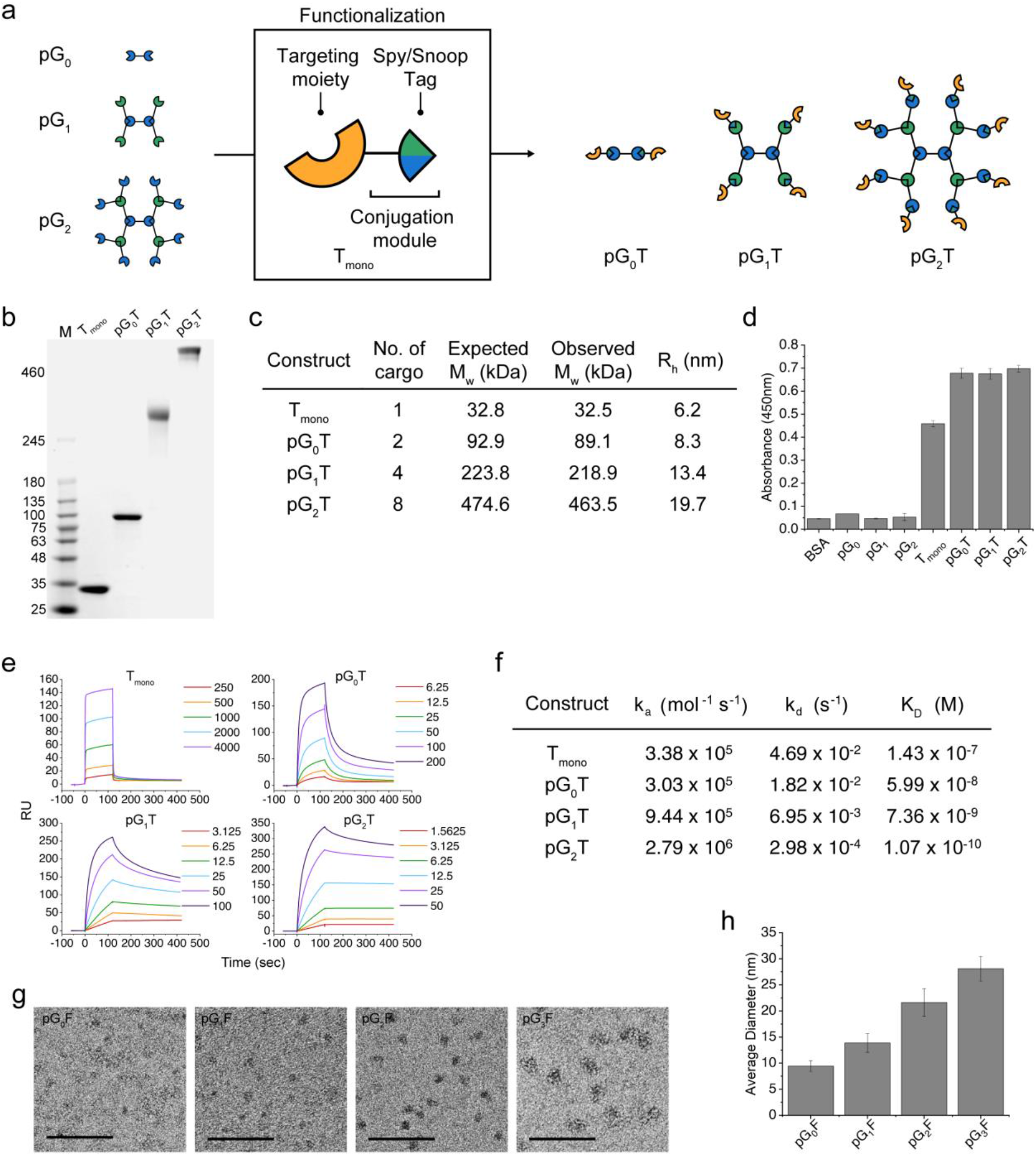
Functionalization of prodrimers using a conjugation module. (**a**) Graphical representation of the functionalization of different prodrimer generations with a targeting moiety. The conjugation module comprises of a tandem of SpyTag and SnoopTag linked using a GS linker. A targeting moiety is genetically fused to the N-terminal of the conjugation module to produce T_mono_. Using either SpyCatcher or SnoopCatcher at the periphery depending on the generation, the prodrimers can be easily functionalized using T_mono_ carrying biomolecules such as a target-specific protein binder, yielding pG_0_T, pG_1_T, and pG_2_T. (**b**) SDS-PAGE (3%–12% gradient) of the functionalized prodrimers with T_mono_ carrying an EGFR-specific repebody as a model targeting moiety. (**c**) Summary of the absolute molecular masses and hydrodynamic radii of the functionalized prodrimers by MALS and DLS, respectively (**Fig. S4b, c, Supporting Information**). (**d**) Direct ELISA of the prodrimers functionalized with T_mono_ carrying the EGFR-specific repebody. Both functionalized and non-functionalized prodrimers were tested. (**e**) SPR sensorgrams of T_mono_ and the functionalized prodrimers with T_mono_ toward human EGFR ectodomain. The legends indicate the concentrations of the prodrimer in nM. (**f**) Summary of association and dissociation rates of the functionalized prodrimers based on the sensograms shown in (e). **(g)** TEM images of different generation prodrimers functionalized with eGFP. The scale bar represents 100 nm. **(h)** Average diameter of the functionalized prodrimers with eGFP shown in (g).

We next checked the binding property of the prodrimers functionalized with T_mono_ carrying an EGFR-specific repebody. As shown in **Figure 2d**, the functionalized prodrimers exhibited distinct signals for human EGFR ectodomain in ELISA, whereas negligible signals were observed from the prodrimers without a targeting moiety. It is likely that the EGFR - specific repebody conjugated to the prodrimers maintains its binding capability for human EGFR ectodomain similar to a free repebody. A protein assembly with multiple valency is expected to provide an enhanced binding capability toward a target mainly through the avidity ^[30–32]^. To test this expectation, we measured the binding affinities of the prodrimers functionalized with T_mono_ carrying the EGFR-specific repebody using surface plasmon resonance (SPR) (**Figure 2e**). As a result, the binding affinity the prodrimers for the EGFR ectodomain was shown to increase by approximately one-order of magnitude with the increasing generation. It is noteworthy that pG_2_T carrying eight repebody molecules showed a K_D_ value of 107 pM, which corresponds to a 1,336-fold increase in the binding affinity compared to a free repebody with a K_D_ value of 143 nM. This result clearly demonstrates a typical feature of multivalent prodrimers in the avidity and cooperativity. The dissociation rates were shown to decrease exponentially with the increasing number of the repebody molecules at the periphery, whereas a linear increase in the association rates was observed (**Figure 2f; Figure S4d, Supporting Information**). Based on the results, it is plausible that the cooperativity and avidity make a major contribution to an increase in the binding affinity of the prodrimers functionalized with an EGFR-specific repebody for the target.

We attempted to analyze the prodrimers using TEM, but failed to obtain clear images due to low contrast. We thus functionalized the prodrimers with an eGFP instead of the repebody, since TEM images of eGFP with high contrast were reported ^[33]^. The prodrimers from a zeroth to a third-generation were functionalized with eGFP using the same method as for the repebody, yielding pG_0_F (indicating a zeroth-generation prodrimer functionalized with a **f** luorescence protein) to pG_3_F, followed by TEM imaging. The resulting prodrimers were observed to be highly homogenous in both SDS-PAGE and size-exclusion chromatography (**Figure S5, Supporting Information**). Interestingly, TEM images of the functionalized prodrimers with eGFP through negative staining showed round dark spots with similar size, despite their highly flexible structures. (**Figure 2g; Figure S6, Supporting Information)**. The average diameter of the functionalized prodrimers was estimated to increase by approximately 6.2 nm with each generation from pG_0_F of 9.4 nm to pG_3_F of 28.0 nm. Considering the linker length and protein size, an incremental increase in the average size of different prodrimer generations coincides well with the approximate theoretical size of a single building block (**Figure 2h; Figure S7, Supporting Information**). Further supported by the TEM analysis, the iterative growth of the prodrimers and their functionalization using the conjugation module seem to be highly specific, leading to the protein assemblies with a well-defined size and valency.

### 2.4. Prodrimers as an intracellular protein delivery platform

From the observation that the prodrimers showed a highly enhanced binding capability due to the avidity, we thought that the prodrimers could be used as a platform for an intracellular protein delivery. For this, a protein delivery system was constructed by combining the prodrimers with our previous work ^[34, 35]^. As depicted in Figure 3a, the protein delivery system is composed of three protein modules which are genetically fused using a flexible linker: a translocation module comprised of a cell-targeting moiety and a translocation domain (TDP) of *Pseudomonas aeruginosa* exotoxin, a cargo module and a conjugation module ^[36]^. As a model cargo, eGFP was employed. An additional KDEL and cathepsin-B cleavage site were introduced into the C-terminal of the cargo for a retrograde translocation and release of the cargo in the endosome, respectively ^[37, 38]^. Lastly, a conjugation module was fused to yield D_mono_ (indicating a monomeric cargo for delivery). The prodrimers functionalized with D_mono_ is expected to deliver a cargo protein to the cytosol as depicted in Figure S8 (Supporting Information). The EGFR-specific repebody targets the cell-surface EGFR, and the functionalized prodrimers undergo receptor-medicated endocytosis. In the endosome, the translocation domain and the cargo is released from the receptor-binding domain and the prodrimer through the cleavage by TDP furin and cathepsin-B, respectively. The cargo is then translocated to the ER by the KDEL receptor, followed by the release to the cytosol. The resulting construct was expressed as a soluble form in *E. coli* and purified, followed by conjugation to pG_0_ and pG_2_, thereby producing pG_0_D to pG_2_D. From a monomeric D_mono_ of 74.3 kDa, the molecular mass of the octomeric pG_2_D increased to 806.6 kDa (**Figure S9a, Supporting Information**). The functionalized prodrimers with D_mono_ were observed to be highly homogeneous, exhibiting single bands on SDS-PAGE, and single peaks in both the size exclusion chromatography and DLS (**Figure 3b; Figure S9b-e, Supporting Information**).

**Figure 3.**
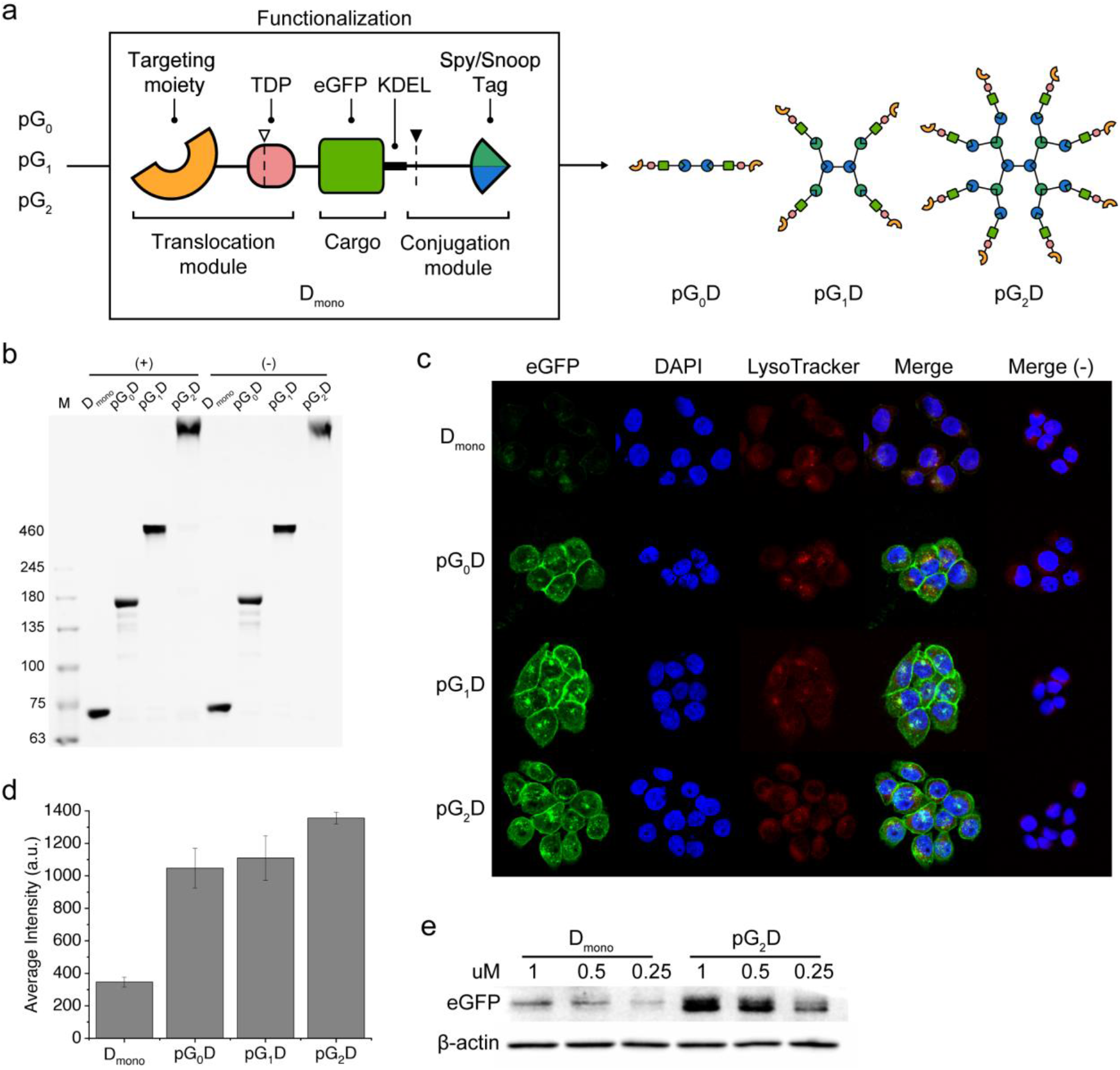
Prodrimer-based intracellular protein delivery. (**a**) Graphical representation of a construct for intracellular protein delivery. The translocation module is composed of a cell-targeting moiety and a translocation domain (TDP) of the exotoxin. A cargo protein is fused to a KDEL sequence, and the cargo and the translocation module are genetically linked to the conjugation module, yielding D_mono_. All protein domains are linked with flexible linkers. As a model, an EGFR-specific protein binder and eGFP were used as a cell-targeting moiety and a cargo, respectively. The dotted line with an empty triangle on the TDP and that with a filled triangle at the end of eGFP represent the furin and cathepsin B cleavage site, respectively. Different generations of prodrimers are functionalized with D_mono_ to produce pG_0_D, pG_1_D, and pG_2_D, respectively. (**b)** SDS-PAGE (3%–6% gradient) of D_mono_ and the functionalized prodrimers. The prodrimers functionalized with D_mono_ carrying an EGFR-specific repebody were designated as (+), whereas those functionalized with D_mono_ carrying an off-target repebody were represented as (-). (**c**) Confocal images of high EGFR-expressing A431 cells after treatment with the functionalized prodrimers for 6 h. Merge (-) indicates the prodrimers functionalized with D_mono_ carrying an off-target repebody. (**d**) Average cell fluorescence intensity of eGFP from A431 cells treated with the same prodrimers as in (c). (**e**) Western blot analysis of intracellular eGFP in lysate of A431 cells treated with the D_mono_ and pG_2_D, respectively. μ_2_;-Actin was used as a control.

We tested the prodrimers functionalized with D_mono_ in terms of the intracellular delivery of the cargo using confocal microscopy. High EGFR-expressing A431 cells were incubated with pG_0_D and pG_2_D for 6 h with a total of 1 μM cargo protein. The functionalized prodrimers have different numbers of cargo proteins depending on the generation. Thus, to match the total concentration of monomeric D_mono_ of 1 μM, the concentrations of the treated pG_0_D, pG_1_D, and pG_2_D were fixed at 500, 250, and 125 nM, respectively. Although the eGFP fluorescence of D_mono_ was difficult to distinguish, the cells treated with pG_2_D showed a distinct fluorescence despite a lower total concentration of treated prodrimer (**Figure 3c**). The average fluorescence intensity of pG_2_D was approximately 3-times greater than that of D_mono_ (**Figure 3d**). The increase in fluorescence intensity can be explained by the enhanced binding affinity of pG_2_D owing to the avidity, as described above. Because pG_2_D has a much lower dissociation rate, a large portion of pG_2_D will be in a bound state in comparison to D_mono_, resulting in a significantly enhanced internalization of a cargo into the cells. The florescence signals were observed to be distributed throughout the cytosol with negligible co-localization with lysotrackers (**Figure S10, Supporting Information**). A similar trend was observed in a western blot analysis of A431 cells (**Figure 3e**). The enhanced cargo delivery by the prodrimers was further examined for another EGFR positive cell line, MDA-MD-468, and a similar result was observed (**Figure S11a, b, Supporting Information**). The prodrimers functionalized with a delivery module carrying an off-target repebody showed negligible fluorescence in both A431 and MDA-MB-468 cells (**Figure S11c, d, Supporting Information**). Similarly, no fluorescence signals were observed in the EGFR-negative cell line, MCF7 (**Figure S12, Supporting Information**). As the results indicate, the prodrimers can be effectively used as an intracellular protein delivery platform, enabling highly efficient protein delivery mainly owing to the avidity.

### 2.5. A cytotoxic protein delivery using the prodrimer

To further assess the utility of the prodrimers as a protein delivery platform, we intended to deliver a cytotoxic protein cargo into cells, and used gelonin as a model cytotoxic protein. Gelonin is a plant-derived N-glycosidase, which inactivates the 60S ribosomal subunit and is known to be unable to cross the cell membrane alone [34, 39]. For this, eGFP was replaced with gelonin in the previous construct, yieldi ng Gmono (indicating a monomeric cargo with gelonin) (Figure 4a). The resulting construct with a size of 75.7 kDa was expressed as a soluble form in *E. coli* and used for functionalization of the prodrimers. From the monomeric Gmono, the octomeric pG_2_G had a molecular mass of 836.8 kDa (Figure S13a, Supporting Information). The functionalized prodrimers were observed to be highly homogeneous, exhibiting single bands on SDS-PAGE (3%–6% gradient), and appeared as clear single peaks in size exclusion chromatography and DLS as well (**Figure 4b; Figure S13b-e, Supporting Information**).

**Figure 4.**
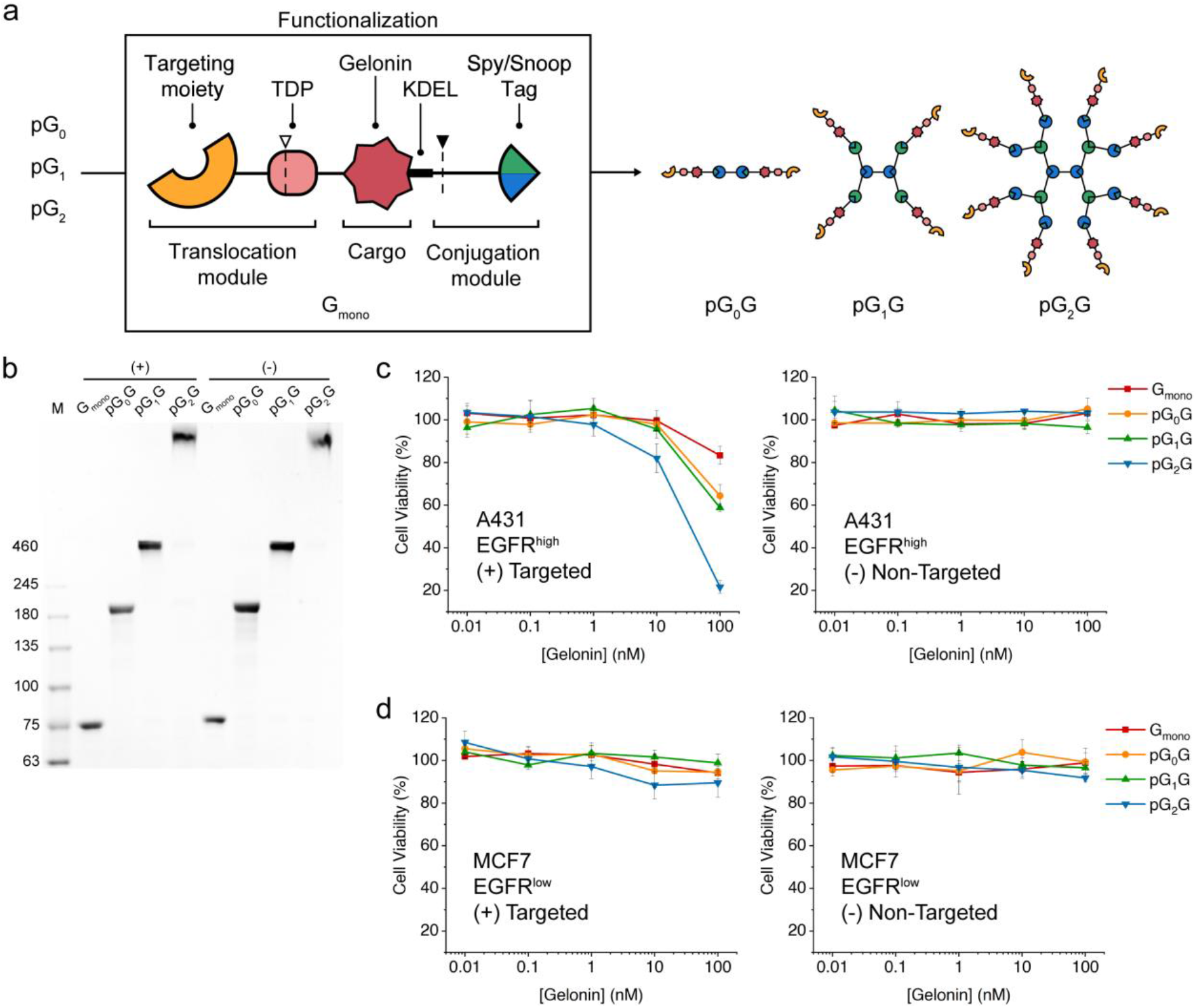
Delivery of a cytotoxic protein cargo by prodrimers. (**a**) Graphical representation of the delivery of gelonin as a cytotoxic protein cargo. Gelonin is shown in red. The dotted line with an empty triangle on the TDP and that with a filled triangle at the end of the gelonin represent the furin and cathepsin B cleavage sites, respectively. Gelonin is genetically fused to the translocation and conjugation modules, yielding Gmono. Different generations of prodrimers are functionalized with Gmono to produce pG_0_G, pG_1_G, and pG_2_G, respectively. (**b**) SDS-PAGE (3%–6% gradient) of the prodrimers functionalized with Gmono carrying either an EGFR-specific repebody (+) or an off-target repebody (-). (**c**) Viability of A431 cells after treatment with the prodrimers functionalized with Gmono carrying either the EGFR-specific repebody (left panel) or an off-target repebody (right panel) for 12 h. **(d)** Viability of MCF-7 cells after treatment with the functionalized prodrimers with Gmono carrying either the EGFR-specific repebody (left panel) or an off-target repebody (right panel) for 12 h.

We investigated the cytotoxic effects of the prodrimers functionalized with Gmono carrying the EGGR-specific repebody against four different cancer cell lines. Two EGFR-positive cell lines (A431 and MDA-MB-468) and two EGFR-negative cell lines (MCF7 and NIH3T3) were tested. To match the total amount of gelonin treated, a lower concentration of the Gmono-functionalized prodrimers was added with the increasing prodrimer generations. As expected, the octomeric pG_2_G showed a much higher cytotoxicity than lower generations of prodrimers even though the total amount of gelonin treated was same. For the A431 cell line, pG_2_G exhibited a remarkable cytotoxic effect, resulting in a cell viability of 21%, whereas other prodrimers resulted in a cell viability ranging from approximately 93% to 60% (**Figure 4c**). A 10-fold higher concentration of monomeric Gmono was required to obtain the same cytotoxicity as the octomeric pG_2_G **(Figure S14, Supporting Information).** A negligible change in cytotoxicity was observed for the non-functionalized prodrimers (**Figure S15a, Supporting Information**). Similar results were obtained for the MDA-MB-468 cell line (**Fig. S15b, Supporting Information**), while a negligible cytotoxicity was observed in EGFR-low and EGFR-negative cell lines (**Figure 4d; Figure S15c, Supporting Information**). The functionalized prodrimers with Gmono carrying an off-target repebody exhibited no cytotoxic effect on all the tested cell lines (**Figure 4c, d; Figure S15d, e, Supporting Information**). Based on the results, it seems that a significantly increased cytotoxic effect by pG_2_G can be attributed to an enhanced intracellular delivery of a protein cargo mainly owing to the avidity of the prodrimers with multiple valency. Because higher generation prodrimers showed a higher avidity, an increase in the binding affinity of such prodrimers functionalized with a cell-targeting moiety is the most likely reason for the higher cytotoxicity.

## 3. Discussion

The quest for the protein assembly with a tunable size and valency has been increasingly high as a versatile platform for creating novel functional nanostructures with a wide range of applications such as vaccines, drug delivery, biosensors, and disease diagnosis and therapy. We demonstrated an efficient and simple bottom-up approach to constructing a supramolecular protein assembly in a way that allows the size and valency to be controlled using only a core protein and two monomeric protein building blocks. Using an orthogonal system paring both SpyCatcher/Spytag and SnoopCatcher/SnoopTag, a dendrimer-like protein assembly, namely prodrimer, was grown to a mega-dalton scale in a highly efficient and stepwise manner through an iterative and alternate addition of a building block. The largest protein assembly constructed in this study was the fourth-generation prodrimer, pG_4_, with 950.5 kDa and 20.6 nm, the size of which is comparable to various viruses^[3]^. In principle, a higher generation of prodrimers can be constructed through the exact same procedure. Unlike the protein assemblies found in nature, the present approach allows the use of diverse building blocks with different sizes and structures, leading to the protein assembly with high homogeneity in a programmable manner. To the best of our knowledge, this is the first to report a programmable supramolecular protein assembly with a tunable size and valency depending on the generation.

In order for the protein assemblies to be used practically, their functionalization with diverse biomolecules is crucial. As the reactive termini of the prodrimers increases by a factor of 2 with each increase in generation, the valency is scalable accordingly. This allows for an easy functionalization with a controllable number of biomolecules through the use of a conjugation module composed of a tandem of SpyTag and SnoopTag. While a linear chain of proteins was constructed using the orthogonal Spy/SnoopCatcher, such linear assembly has some limitations as a protein assembly since a single growth step is required for every cargo extension ^[26]^. In contrast, the prodrimers can be decorated with diverse number of cargos in a single and efficient way at the periphery, generating protein assemblies with multimodality. When functionalized with a target-specific protein binder, the binding affinity of the prodrimers could be modulated and enhanced through the avidity effect. Specifically, the second-generation prodrimer functionalized with an EGFR-specific repebody exhibited a 1,336-fold increase in the binding affinity for a human EGFR compared to a monomeric equivalent, confirming the effect of the avidity. It is noteworthy that a prodrimer with a higher valency led to an exponential decrease in the dissociation rate, mainly contributing to a high binding affinity for the target. A higher generation prodrimer is expected to show a further enhancement in the binding affinity. Similarly, the second-generation prodrimers functionalized with a delivery module and a protein cargo were shown to have a significantly enhanced intracellular translocation of the protein cargo compared to a monomeric equivalent mainly owing to the avidity. When gelonin was used as a model cytotoxic cargo, the cytotoxic effect on the cell viability was significantly enhanced, even though the same total concentration of gelonin was used. This result exemplifies the utility of the prodrimers as an intracellular protein delivery platform. The controllability of the size and valency of prodrimers is also an advantage in *in vivo* environments because a higher target binding affinity and a longer blood circulation time of the prodrimers will lead to a higher therapeutic efficacy. Furthermore, the prodrimers are biodegradable and offer a high biocompatibility compared to inorganic nanomaterials.

## 4. Conclusion

In conclusion, we demonstrated the programmable growth and utility of a dendrimer-like supramolecular assembly of proteins, called “prodrimers,” as a novel biocompatible platform for various applications such as vaccines, drug delivery, biosensors, and disease diagnosis and therapy. The supramolecular protein assembly was efficiently constructed through a simple bottom-up approach in a way that allows the size and valency to be controlled using only a core protein and two monomeric protein building blocks. The prodrimers with sizes ranging from small proteins to virus capsids were constructed through a stepwise and iterative growth using a core protein and two building blocks in a programmable manner. Furthermore, the functionalization of the prodrimers was easy to achieve through a simple process using a conjugation module. The multivalent nature of the prodrimers showed a distinct avidity effect, leading to a significantly increased binding affinity for a target compared to a monomeric equivalent. Consequently, the prodrimers functionalized with a cytotoxic protein cargo exhibited a higher cytotoxic effect on the cell viability. Taken together, the present approach can be effectively used in the creation of advanced protein assemblies for biotechnological and medical applications.

---

**Scheme 1.**
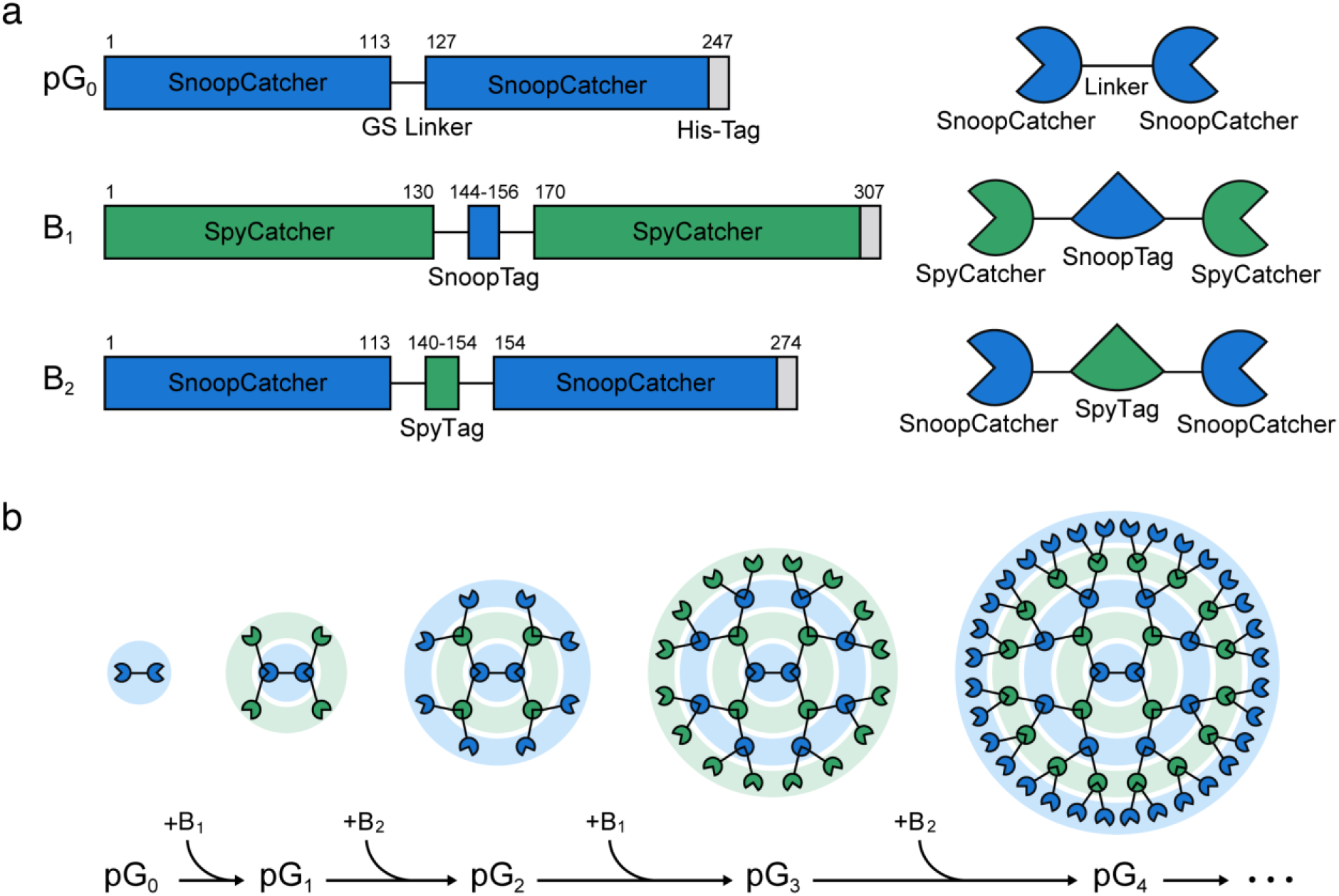
Schematic of the assembly of proteins through iterative and alternate additions of a building block protein. (**a**) Graphical representation of three monomer proteins used for the construction of the protein assembly, termed prodrimers: the core protein (pG_0_) and two building block proteins (B_1_ and B_2_). The numbers indicate the lengths of the amino acids. Blue and green colors represent orthogonal pairs of SpyCatcher/SpyTag and SnoopCatcher/SnoopTag, respectively. The core protein (pG_0_) comprises a tandem of SnoopCatcher linked to each other using a GS linker. The two building blocks (B_1_ and B_2_) are composed of two tandem repeats of SpyCatcher and SnoopCatcher linked with SnoopTag and SpyTag, respectively. (**b**) Stepwise growth of prodrimers through an iterative and alternate addition of a building block protein. Starting from the core protein pG_0_, an alternate addition of a building block protein (B_1_ or B_2_) leads to a sequential growth in the size and valency of prodrimers.

