## Supplementary Information for "Dendrimer-like supramolecular assembly of proteins with a tunable size and valency through stepwise iterative growth"

**Experimental Section**

**Protein expression and purification**

All genes encoding the proteins used in the present work were cloned into a pET21a vector (Novagen) using NdeI and XhoI restriction sites. The genes for SpyCatcher and SnoopCatcher were synthesized as a gBlock Gene Fragment from Integrated DNA Technologies (IDT). SpyTag and SnoopTag were amplified using polymerase chain reaction. The protein monomers were linked to each other with a flexible linker (GSAGSAAGSGEF) ^[1]^. For the construction of an intercellular protein delivery module, the gene coding for an EGFR-targeting repebody, an off-target repebody, the translocation domain of *Pseudomonas aeruginosa* exotoxin, gelonin, and eGFP were amplified using PCR. Each protein domain was connected using a flexible GGGS linker, and a 6xHis-Tag was fused to the C-terminal of each construct. Amino acid sequences of proteins used for the construction and functionalization of protein assemblies are shown (**Table S1,** S**2**, **Supporting Information**) ^[2-4]^. For protein expression, the vectors harboring the gene were transformed into *E. coli* BL21 (DE3) and grown at 37 °C. When the optical density reached 0.6–0.8 at 600 nm, IPTG (0.5 mM) was added for induction. The induced cells were further grown for 18 h at 18 °C and harvested through centrifugation at 8000 rpm.

For protein purification, the harvested cells were resuspended in a lysis buffer (50 mM Tris, 300 mM NaCl, and 10 mM Imidazole at pH 7.8) and disrupted through sonication. Following centrifugation at 18,000 rpm for 1 h, the supernatant was loaded into a Ni-NTA Superflow (Qiagen). The loaded Ni-NTA column was washed with a wash buffer (50 mM Tris, 300 mM NaCl, and 20 mM Imidazole at pH 7.8). Finally, the proteins were eluted with an elution buffer (50 mM Tris, 300 mM NaCl, and 250 mM Imidazole at pH 7.8). All eluted proteins were further purified through size exclusion chromatography (Superdex 200, GE Healthcare). The monomers for protein assembly were purified using a sodium borate buffer (50 mM at pH 10.5), and constructs of gelonin and eGFP cargo were purified using a 20 mM Tris buffer (300 mM NaCl at pH 7.8).

**Cell Culture**

A431 (human epidermoid carcinoma, ATCC No. CRL-1555) and MCF7 (human adenocarcinoma, ATCC No. HTB-22) were cultivated in an RPMI medium (Capricorn) supplemented with 10% (v/v) fetal bovine serum (FBS, Hyclone). MDA-MB-468 (human adenocarcinoma, ATCC No. HTB-132) and NIH3T3 (mouse embryonic fibroblast, ATCC No. CRL-1658) were cultivated in DMEM (Capricorn) supplemented with 10% (v/v) FBS. All cell cultures were carried out in a 5% CO_2_ chamber at 37 °C.

**Construction and purification of protein assembly**

For the protein assembly, the core protein (pG_0,_ zeroth-generation prodrimer) was incubated with an excess of 2–4 molar ratio equivalent of the total available termini using a building block protein in 50 mM sodium borate buffer (pH 9.0, 0.1% Tween-20). The resulting first-generation prodrimer (pG_1_) was purified using size exclusion chromatography (Superdex 200, GE Healthcare) in a 50 mM sodium borate buffer (pH 10.5). The fractions for pG_1_ were collected and concentrated using Amicon Ultra-15 Centrifugal Filters (Merck). For a further growth of the protein assembly, the same step was repeated using an alternate building block protein, yielding a prodrimer with a higher generation. For functionalization with protein cargos, each generation prodrimers were incubated using 3 molar ratio equivalent to the total available termini using a protein cargo which had been genetically fused to a conjugation module comprising a tandem of SpyTag and SnoopTag in a 20 mM Tris buffer (300 mM NaCl at pH 7.8) with 0.1% Tween-20. The functionalized prodrimers were purified using size exclusion chromatography (Superdex 200, GE Healthcare) in a 20 mM Tris buffer (300 mM NaCl at pH 7.8). For the analysis of the assembly and homogeneity of prodrimers, a Superdex Increase 200 10/300 (GE Healthcare) column was applied.

**Confocal microscopy**

For confocal imaging of the cells treated with prodrimers, the respective cells were attached to a microscopy 8-well chamber slide (SPL) at a density of 2 × 10^3^ cells/well. After 3 days of growth, the serum-containing medium was treated with an appropriate amount of prodrimers at each generations for 6 h. To track the lysosome and acidic compartments, 200 nM of Lysotracker Red DND-99 (Thermo Scientific) was treated for 2 h. The cells were washed with DPBS containing 0.1% Tween-20 and fixed with 4% (v/v) paraformaldehyde for 20 min. Cell nuclei were counterstained with DAPI in the mounting medium (Vector). Confocal images were obtained using a Zeiss LSM 780 confocal microscope with a ×40 Apochromat objective with a 1.0 numerical aperture (Carl Zeiss).

***In vitro* cytotoxicity**

Cells were seeded in a 96-well plate (SPL) at a density of 5 × 10^3^ cells/well in a medium containing FBS. After growth overnight, the medium was changed to a fresh medium without serum, and an appropriate concentration of prodrimers carrying a protein cargo were added. After incubation for 12 h, the medium was replaced with a fresh medium without serum followed by further incubation for 3 days. Each well was supplemented with 10 μL of a CCK-8 reagent (Dojindo) and incubated for 2 h at 37 °C. Cell viability was determined using an Infinite M200 plate reader (Tecan) by measuring the OD_450_.

**Western blot assay**

The cells were seeded in a 24 well plate (SPL) at a density of 2 × 10^5^ cells/well and grown overnight. The cells were then treated with an appropriate concentration of prodrimers for 6 h, followed by three washes with cold DBPS. The cells were homogenized in a RIPA Lysis and Extraction Buffer (Thermo Scientific) with mild agitation for 30 min at 4 °C. The cell suspension was centrifuged for 20 min at 12,000 rpm, and the supernatant was collected. A total of 80 μg of protein was loaded and separated using a 12% SDS-PAGE gel. The proteins on the gel were transferred to a nitrocellulose membrane (Bio-Rad) at 100 V for 2 h in ice. The membrane was first blocked with PBS containing 0.1% Tween-20 and 3% (v/v) bovine serum albumin (BSA). Anti-GFP monoclonal (Santa Cruz) and anti-beta-actin monoclonal (Santa Cruz) primary antibodies were treated at 4 °C overnight, followed by incubation with HRP-conjugated anti-mouse IgG (Bio-Rad) secondary antibody at 37 °C for 1 h. The membrane was washed with PBST three times after each step and visualized using an enhanced chemi-luminescence solution (Millipore) with ChemiDoc XRS+ (Bio-Rad).

**Dynamic light scattering (DLS)**

Prodrimers were diluted in a PBS buffer (pH 7.5) and subjected to dynamic light scattering (DLS) to measure the hydrodynamic radii using a Zetasizer nano zs (Malvern). All measurements were taken at 25 °C.

**Binding kinetics using surface plasmon resonance (SPR)**

The association and dissociation kinetics of the functionalized prodrimers with an EGFR-targeting protein binder were determined through SPR using a BIACORE 3000 (Biacore AB). Human EGFR ectodomain with an hFc Tag (Sino Biological) was immobilized on a CM5 sensor chip through EDC/NHS conjugation chemistry. A mixture of 0.4 M 1-(3-dimethylaminopropyl)-3-ethylcarbodiimide hydrochloride (EDC) and 0.1M N-hydroxysulfosuccinimide (sulfo-NHS) was used to activate the chip surface, and 20 μg/mL of a human EGFR ectodomain in a sodium acetate buffer (10 mM at pH 4.5) was injected for immobilization onto the chip. Ethanolamine (1.0 M at pH 8.0) was used to block the chip. The functionalized prodrimers were injected at different concentrations. The sensorgrams were fit using a 1:1 Langmuir binding model using the software provided, and the association and disassociation rates were determined.

**Multi-angle light scattering (MALS)**

The absolute molecular masses of proteins were determined using multi-angle light scattering (MALS) (Dawn Heleos II, Enhanced Optical Signals) coupled to a Superdex 200 Increase 10/300 column (GE Healthcare). The absolute molecular masses were determined using the ASTRA program (Wyatt Technologies).

**Transmission electron microscopy (TEM)**

For negative stain TEM imaging, prodrimer samples (0.05mg/ml) were applied to glow-discharged carbon grids and incubated for 1 min. The grids were then washed twice with water and incubated with 1.5% (w/v) uranyl acetate for 1 min. The grids were blotted using filter paper and dried under ambient conditions. The prepared grids were analyzed using a Tecnai F20 200-kV field-emission transmission electron microscope (FEI) with a CCD camera (Gatan).

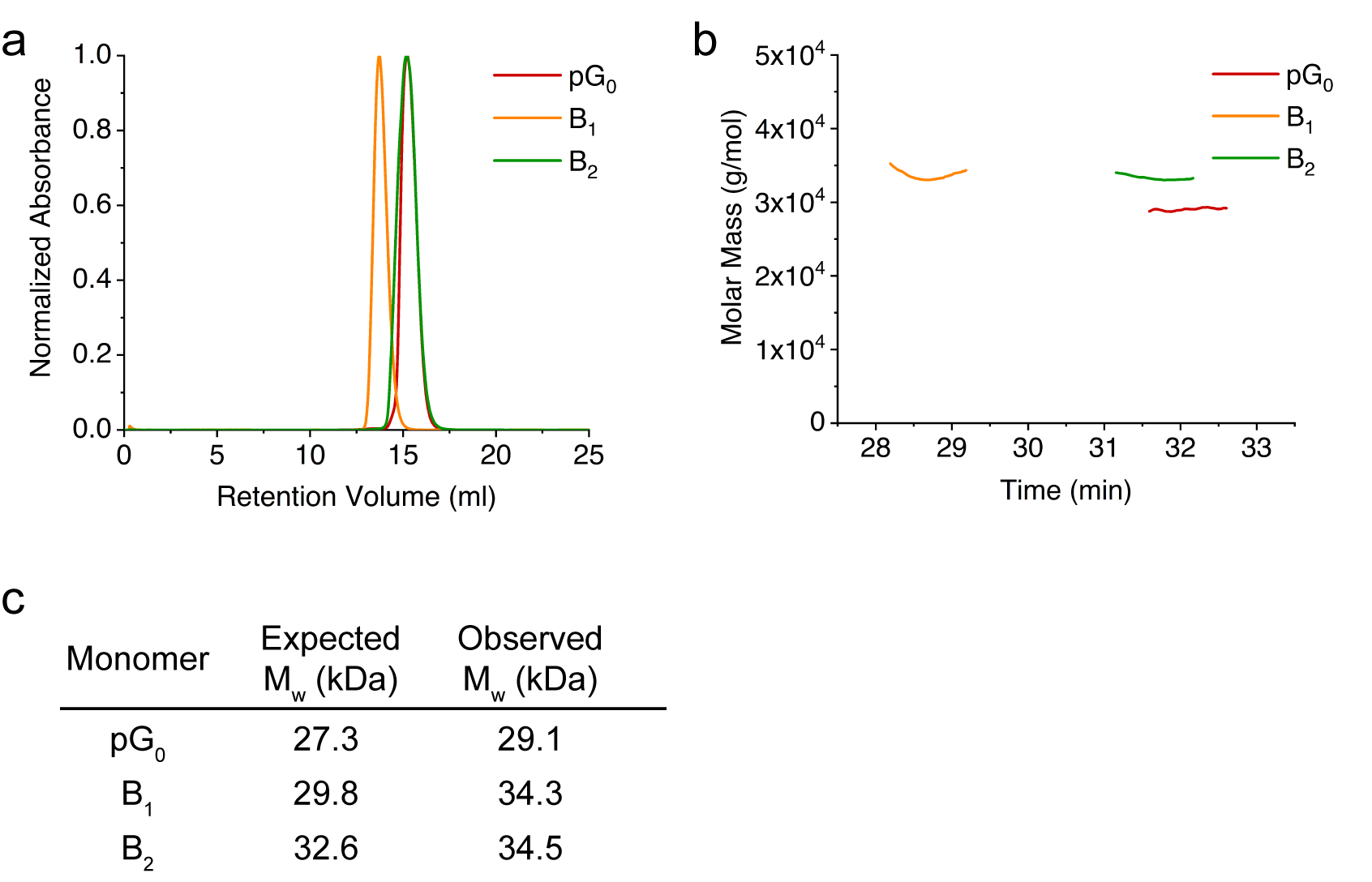

**Figure S1. Purification and biophysical characterization of the core protein and two building blocks for the construction of prodrimers. (a)** Size exclusion chromatography of the core protein (pG_0_) and two building blocks (B_1_ and B_2_). The peaks represent the normalized absorbance of each protein at 280 nm. **(b)** Absolute molecular masses of the core protein and two building blocks by MALS. Eluted protein fractions from SEC were analyzed. **(c)** Summary of the absolute molecular masses of the core and two building blocks determined by MALS in (b).

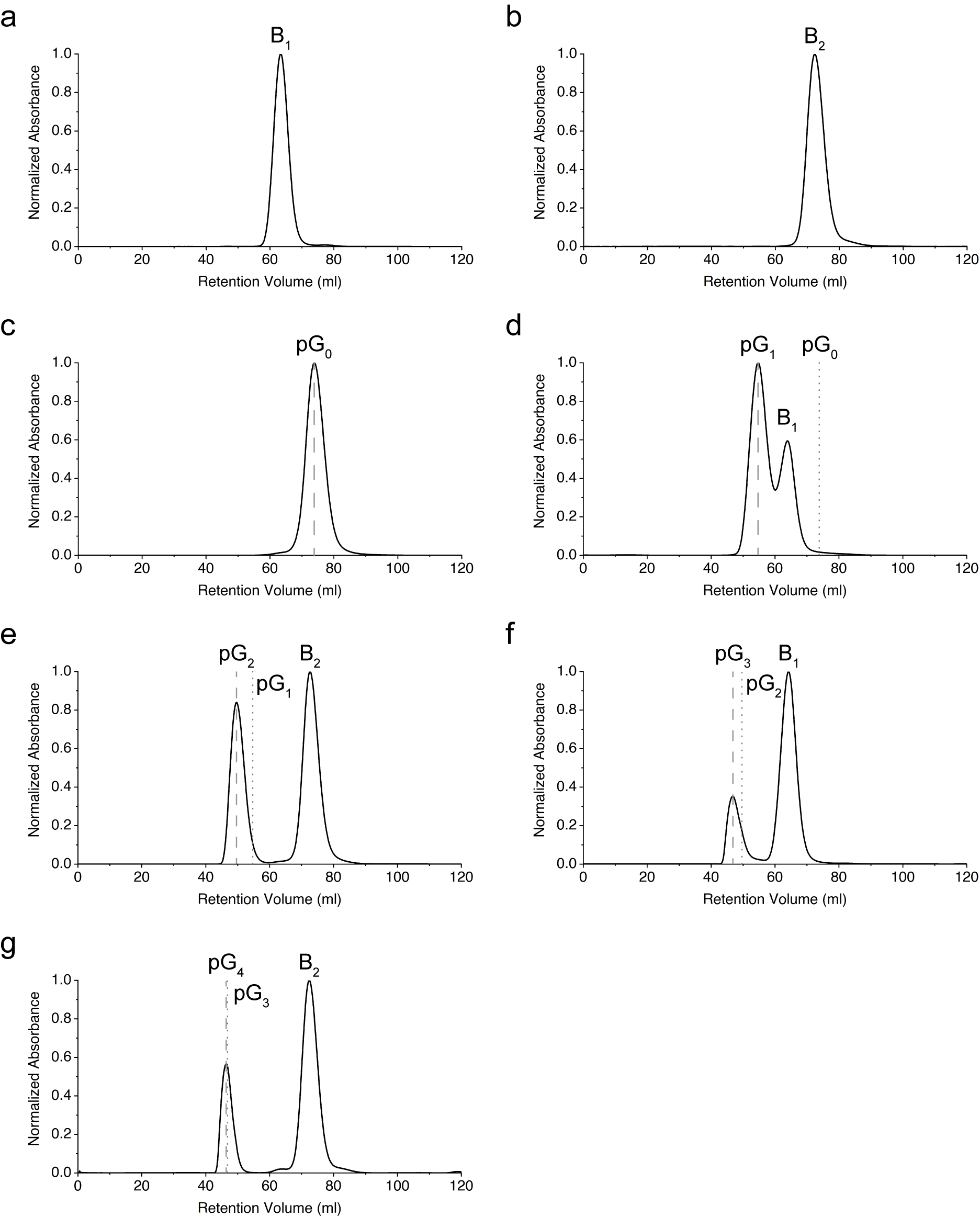

**Figure S2. Construction of different prodrimer generations through a stepwise iterative growth.** Different generations of prodrimers were purified by size exclusion chromatography (SEC) using a S200 column in sodium borate buffer (pH 10.5). The purified prodrimers were used for the construction of the next generation prodrimers. Elution peaks were monitored at 280 nm. **(a)** Building block protein B_1_. **(b)** Building block protein B_2_. **(c)** Zeroth-generation prodrimer pG_0_ (Core protein). **(d)** Construction and analysis of the first-generation prodrimer pG_1_. The core protein pG_0_ was incubated with an excess of a building block protein B_1_ overnight, followed by purification through SEC. The dotted line represents the maximum peak of the previous generation prodrimer (in this case pG_0_), and the dashed line indicates the peak maximum of a next-generation prodrimer (in this case pG_1_). All of pG_0_ were efficiently grown to pG_1_ by addition of B_2_, leaving a negligible trace of the pG_0_ peak. Eluted fractions of pG_1_ were collected for further growth. (**e**) Construction and analysis of the second-generation prodrimer pG_2._ The growth procedure and analysis conditions were the same as in (d). **(f)** Construction and analysis of the third-generation prodrimer pG_3._ **(g)** Construction and analysis of the fourth-generation prodrimer pG_4_. In the case of pG_3_ and pG_4,_ both proteins were eluted near the void fractions due to their large molecular masses, resulting in the overlap of the peak maximum.

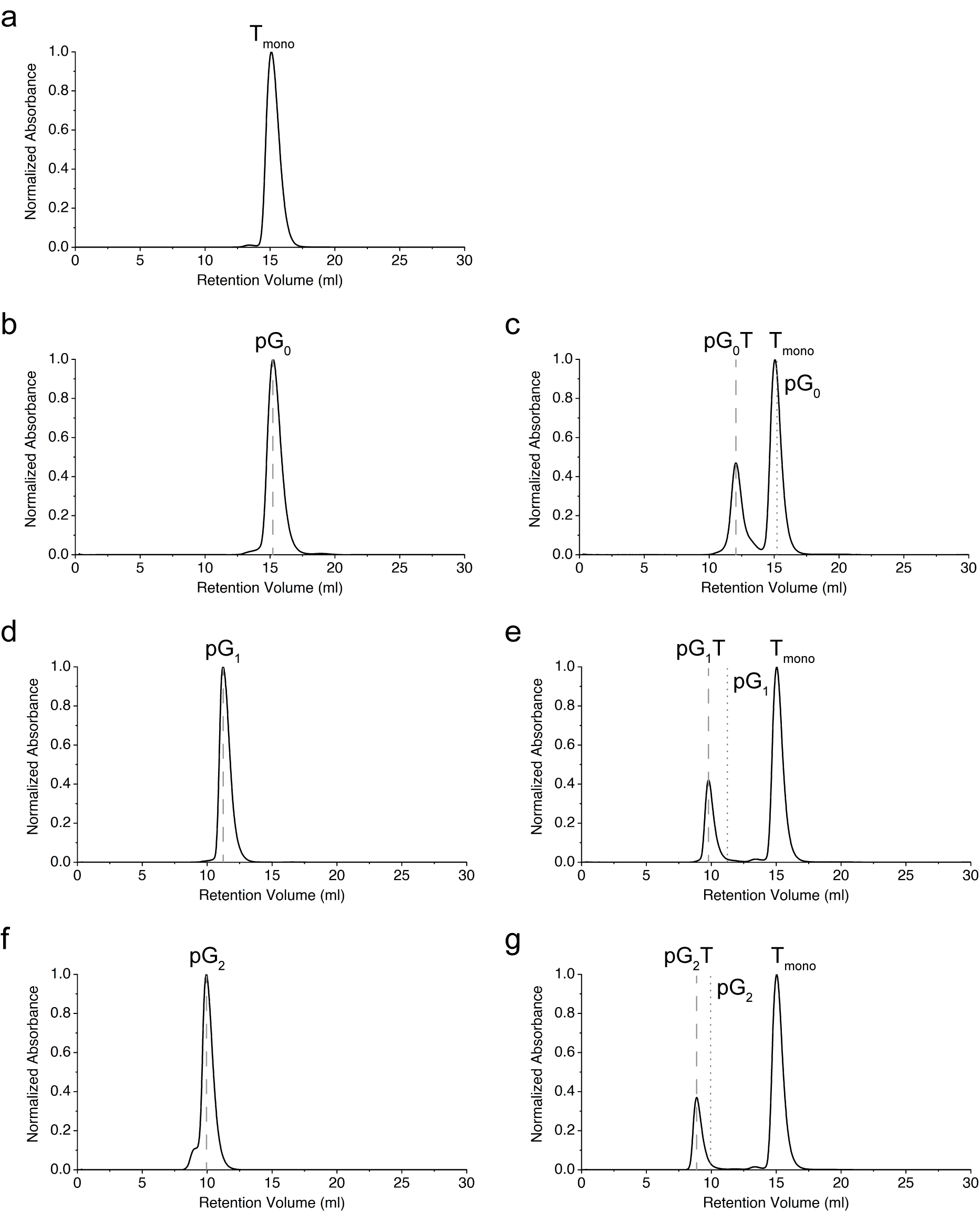

**Figure S3. Functionalization of prodrimers with a targeting moiety and analysis of the functionalized prodrimers through size exclusion chromatography.** The peaks in SEC show the normalized absorbance of the functionalized prodrimers at different generations at 280 nm using a S200 column in Tris buffer (pH 7.5). Void volume is 8.48 ml. **(a)** Purification of an EGFR-specific repebody fused to a conjugation module, T_mono_. The conjugation module comprises of a tandem of SpyTag and SnoopTag linked to each other using a GS linker. **(b)** Zeroth-generation prodrimer pG_0_. **(c)** Formation of pG_0_T by incubating pG_0_ with an excess of T_mono_ overnight. The dotted line represents the peak maximum of the starting prodrimer (in this case pG_0_), and the dashed line indicates the peak maximum of the newly functionalized prodrimer (in this case pG_0_T). (**d**) Purified pG_1._ **(e)** Formation of pG_1_T by incubating pG_1_ with an excess of T_mono_ overnight. **(f)** Purified pG_2_. **(g)** Formation of pG_2_T by incubating pG_2_ with an excess of T_mono_ overnight.

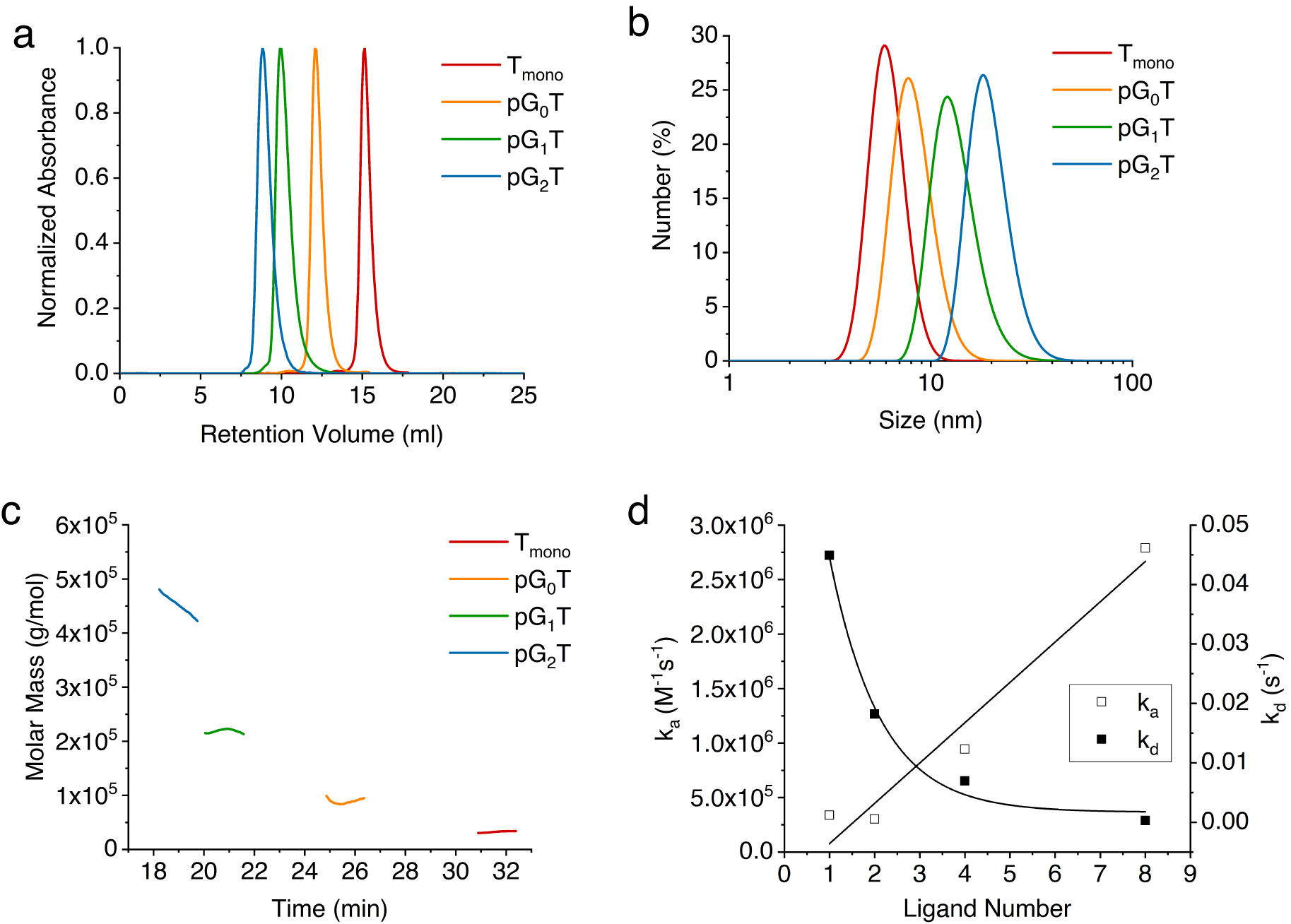

**Figure S4. Biophysical characterization of the prodrimers functionalized with a target-specific protein binder. (a)** Size exclusion chromatography of T_mono_ and different generation prodrimers functionalized with an EGFR-specific repebody. The peaks represent the normalized absorbance of each generation prodrimers at 280 nm. **(b)** Dynamic light scattering of the prodrimers functionalized with the repebody. **(c)** Absolute molecular masses of the prodrimers functionalized with the repebody by MALS. The prodrimers eluted from SEC were analyzed. **(d)** Association and dissociation rates of the functionalized prodrimers based on the sensograms represented in **Figure 2e** as a function of the number of the EGFR-specific repebody molecules.

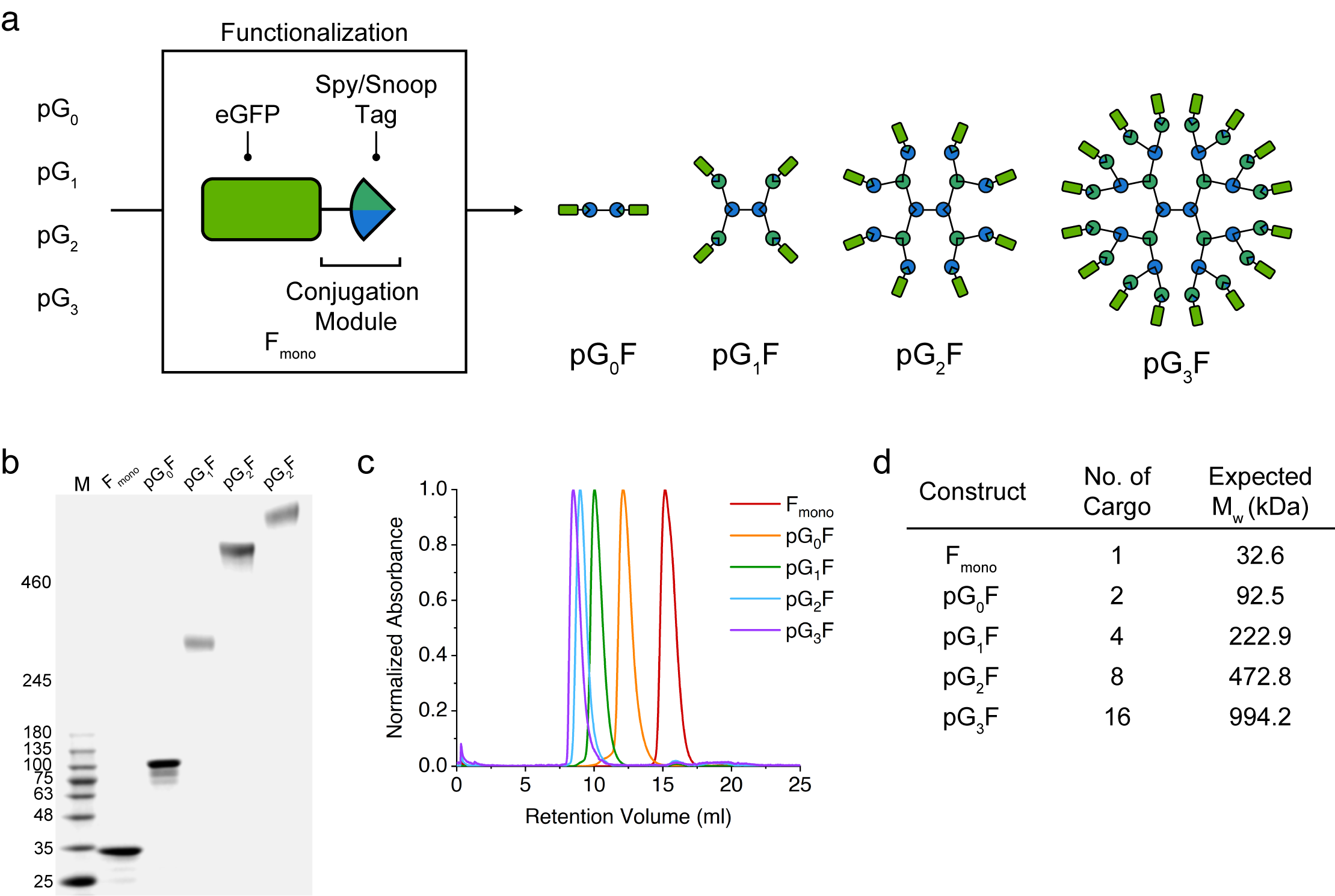

**Figure S5. Functionalization of prodrimers with eGFP and biophysical characterization of the functionalized prodrimers.** **(a)** Graphical representation of the functionalization of different generation prodrimers with eGFP which had been genetically fused to the N-terminal of a conjugation module to produce F_mono_. The conjugation module comprises of a tandem of SpyTag and SnoopTag linked to each other using a GS linker. Using either SpyCatcher or SnoopCatcher at the periphery depending on the generation, the prodrimers were functionalized with eGFP, yielding pG_0_F, pG_1_F, and pG_2_F and pG_3_F. **(b)** SDS-PAGE (3%–12% gradient) of the prodrimers functionalized with eGFP. **(c)** Size exclusion chromatography of the functionalized prodrimers with eGFP. The elution peaks represent the normalized absorbance of the prodrimers at 280 nm. **(d)** Summary of the number of eGFP molecules at different prodrimer generations and their expected molecular masses.

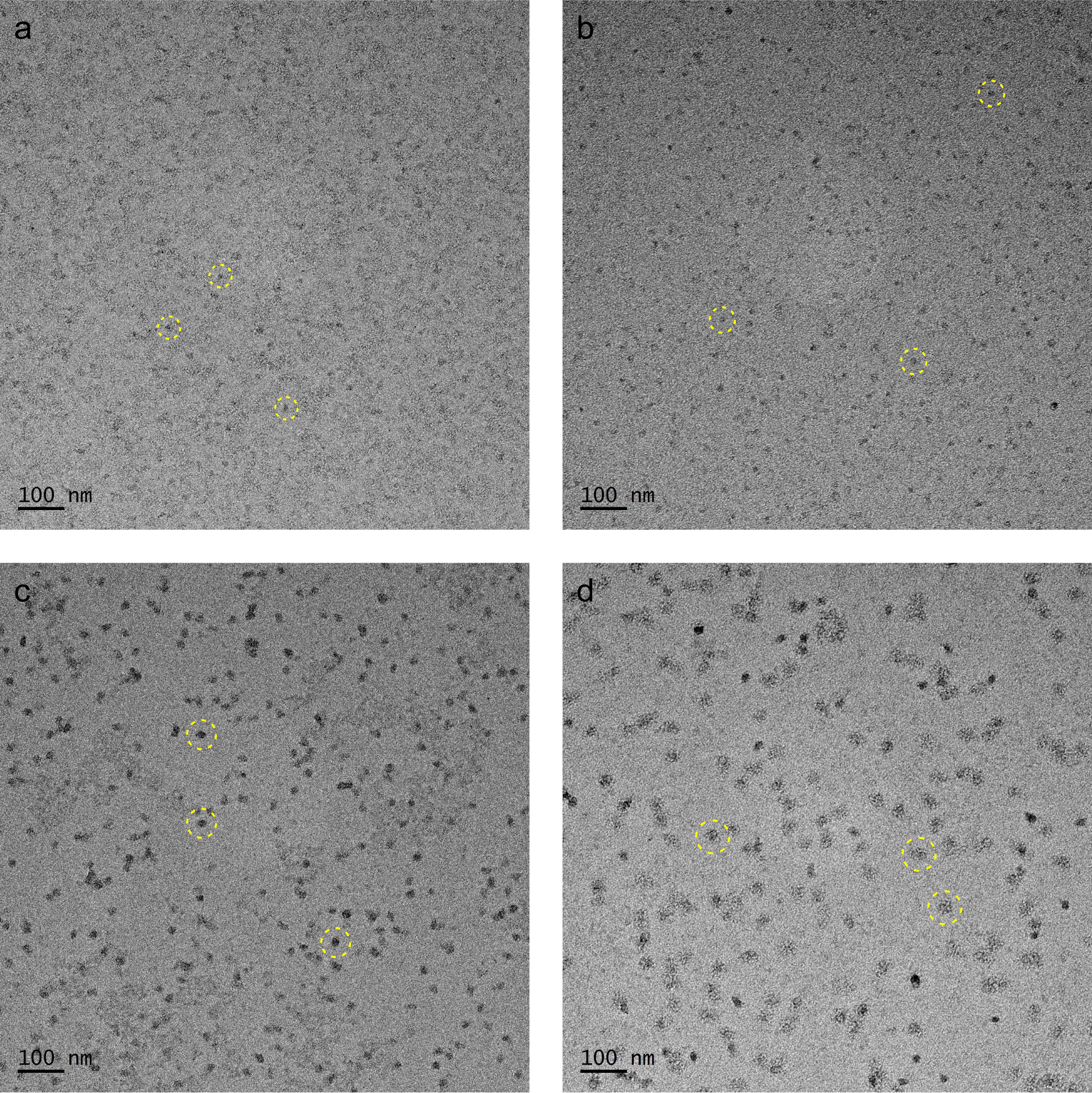

**Figure S6. TEM image of the prodrimers functionalized with eGFP.** Different generations of prodrimers were functionalized with eGFP and subjected to TEM analysis**. (a)** pG_0_F. **(b)** pG_1_F. **(c)** pG_2_F. **(d)** pG_3_F. The images in yellow dashed circles indicate representative prodrimers.

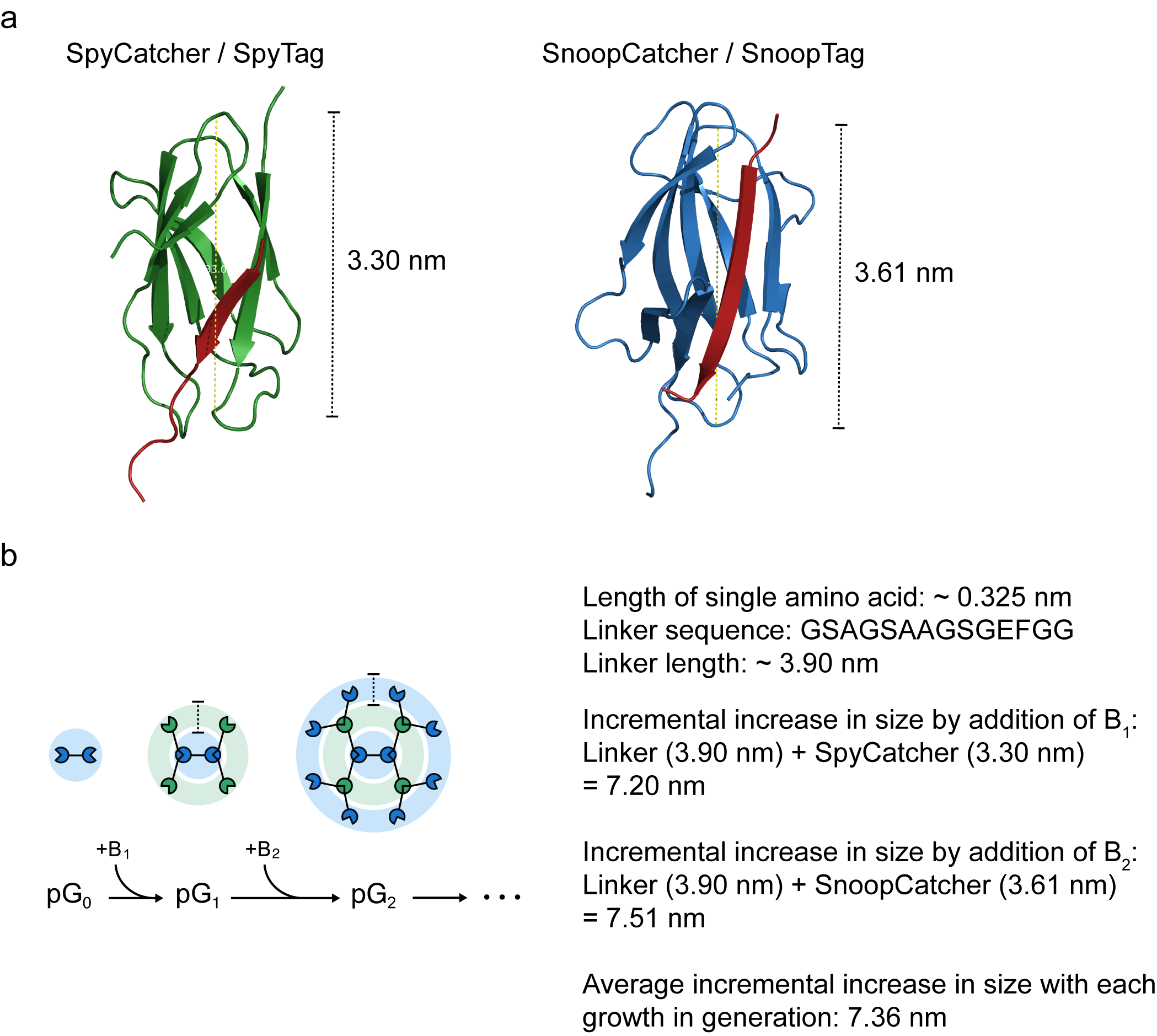

**Figure S7. Approximate incremental increase in the size of prodrimers with the increasing generation. (a)** Structure and size of a SpyCatcher/SpyTag (PDB ID: 4MLI) and a SnoopCatcher/ SnoopTag ((PDB ID: 2WW8) represented by PyMOL. SpyCatcher and the reconstituted SpyTag are indicated by green and red, respectively. SnoopCatcher and the reconstituted SnoopTag are represented by blue and red, respectively. **(b)** According to Scheme 1, the SpyTag/SnoopTag at the middle of a building block is linked to the previous generation prodrimer. Approximate length of a linker used between the each building block protein, given that each amino acid is 0.325 nm and the linker is fully extended ^[5]^. Thus, the increase in the average size in the next generation prodrimer corresponds to the sum of the size of a linker and a single SpyCatcher/SnoopCatcher protein. The maximum growth in the size of prodrimer might be an average 7.36 nm, assuming that the growth is one-dimensional, The theoretical increase in the size of prodrimer with the generation matches the incremental increase of 6.2 nm which was estimated using TEM in **Figure 2h**, considering that the protein in solution are three-dimensional and not completely extended.

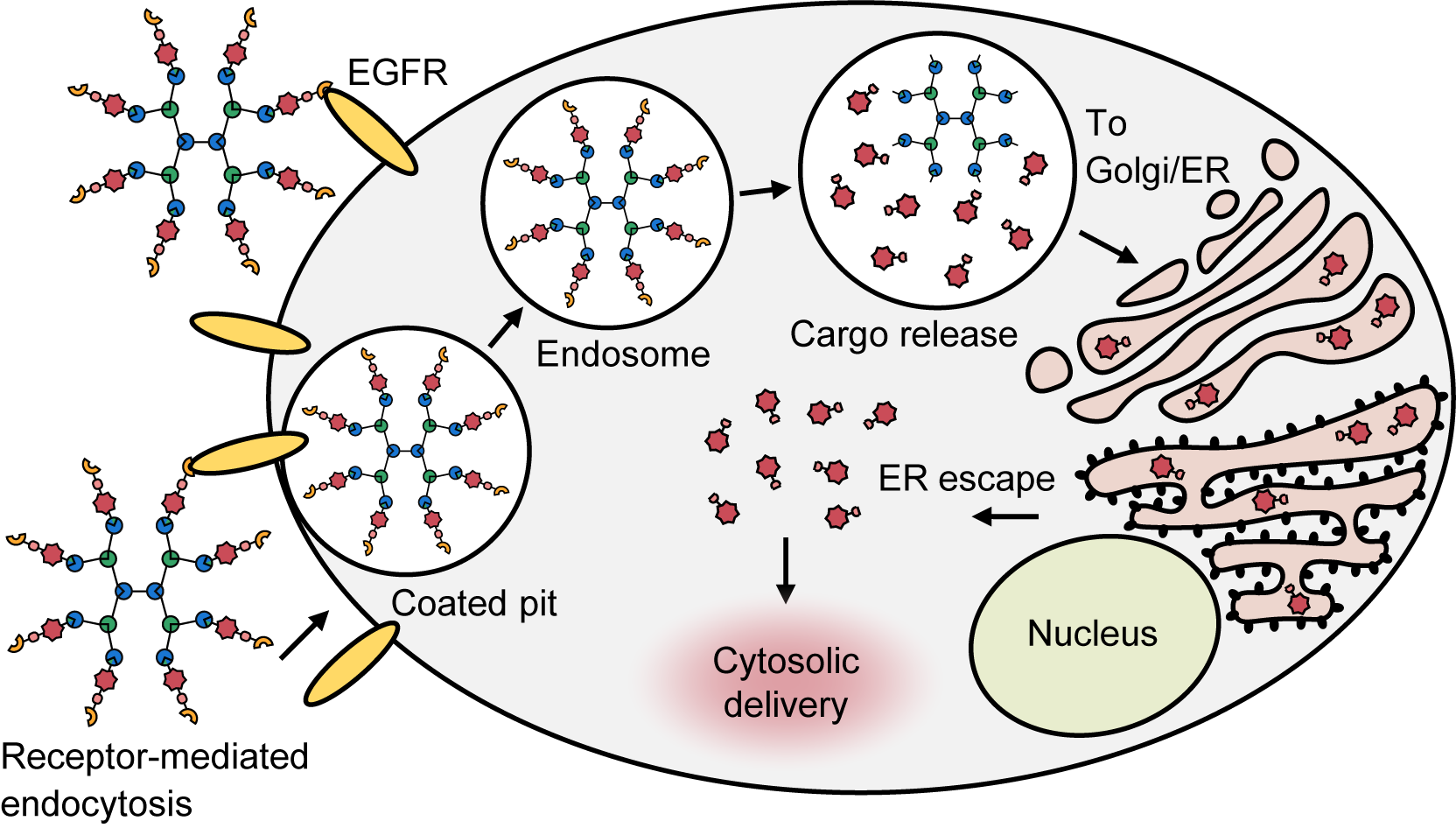

**Figure S8. Scheme of intracellular protein delivery using prodrimers.** A protein cargo was delivered to the cytosol by the prodrimers functionalized with a targeting moiety and a protein cargo. An EGFR-specific repebody on the prodrimer binds the cell surface EGFR, and the functionalized prodrimer undergoes endocytosis. In the endosome, the translocation domain is released from the receptor-binding domain through the cleavage of TDP by furin and cathepsin. The cargo is then translocated to the ER by the KDEL receptor, followed by the release to the cytosol.

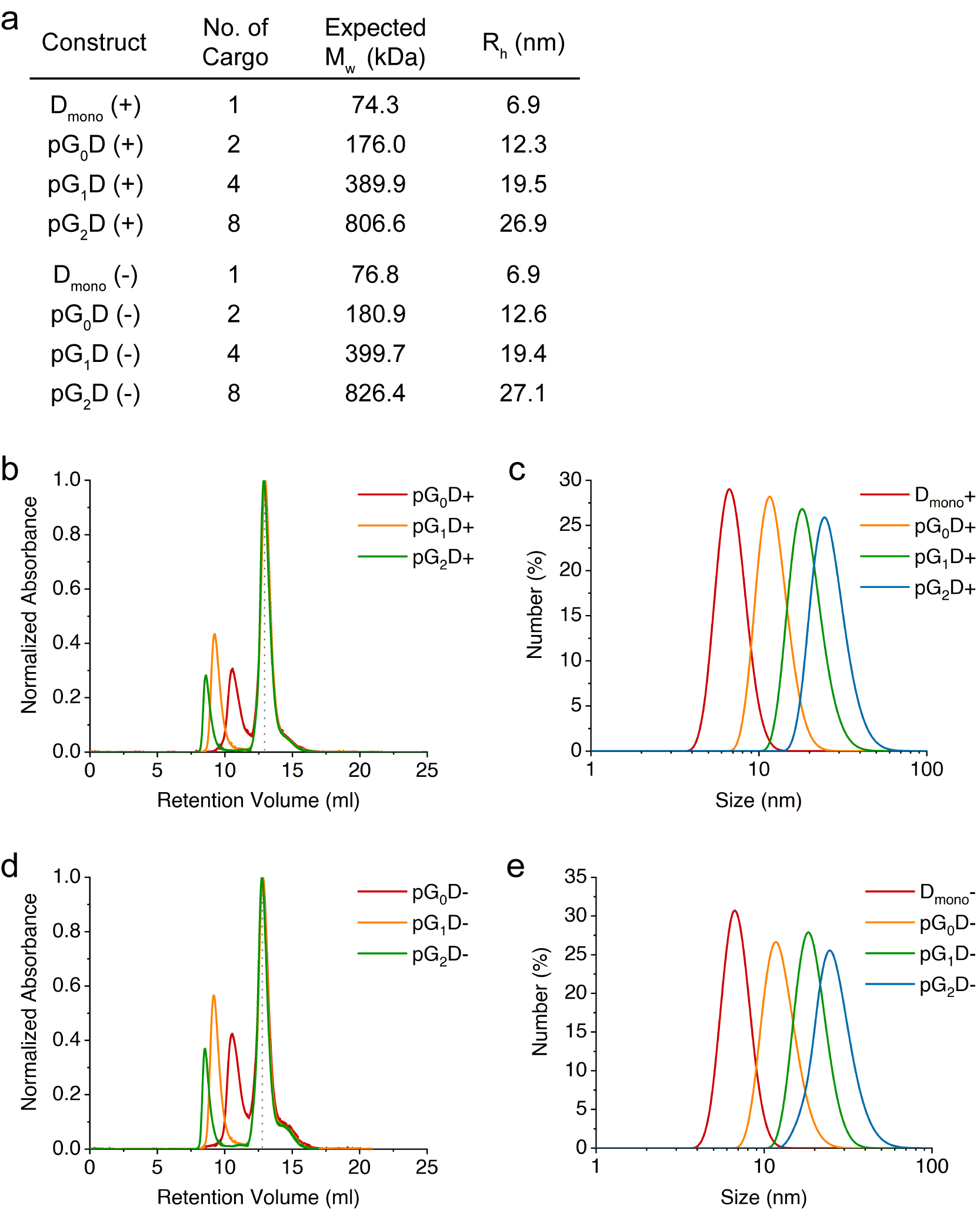

**Figure S9. Biophysical characterization of the prodrimers functionalized with D_mono_. (a)** Summary of the expected molecular masses and the hydrodynamic radii of different prodrimer generations functionalized with D_mono_. (+) and (-) indicate the translocation module carrying an EGFR-specific repebody and an off-target repebody, respectively. (**b**) Size exclusion chromatography of the functionalized prodrimers with the translocation module carrying an EGFR-specific repebody. The peaks represent the normalized absorbance of the prodrimers at 280 nm. The dotted line represent the excess D_mono_ (+) proteins used for functionalization. **(c)** DLS analysis of the prodrimers functionalized with the translocation module carrying an off-target repebody. (**d**) Size exclusion chromatography of the prodrimers functionalized with the translocation module carrying an off-target repebody. The dotted line represent the excess D_mono_ (-) proteins used for functionalization. **(e)** DLS analysis of the prodrimers functionalized with the translocation module carrying an EGFR-specific repebody.

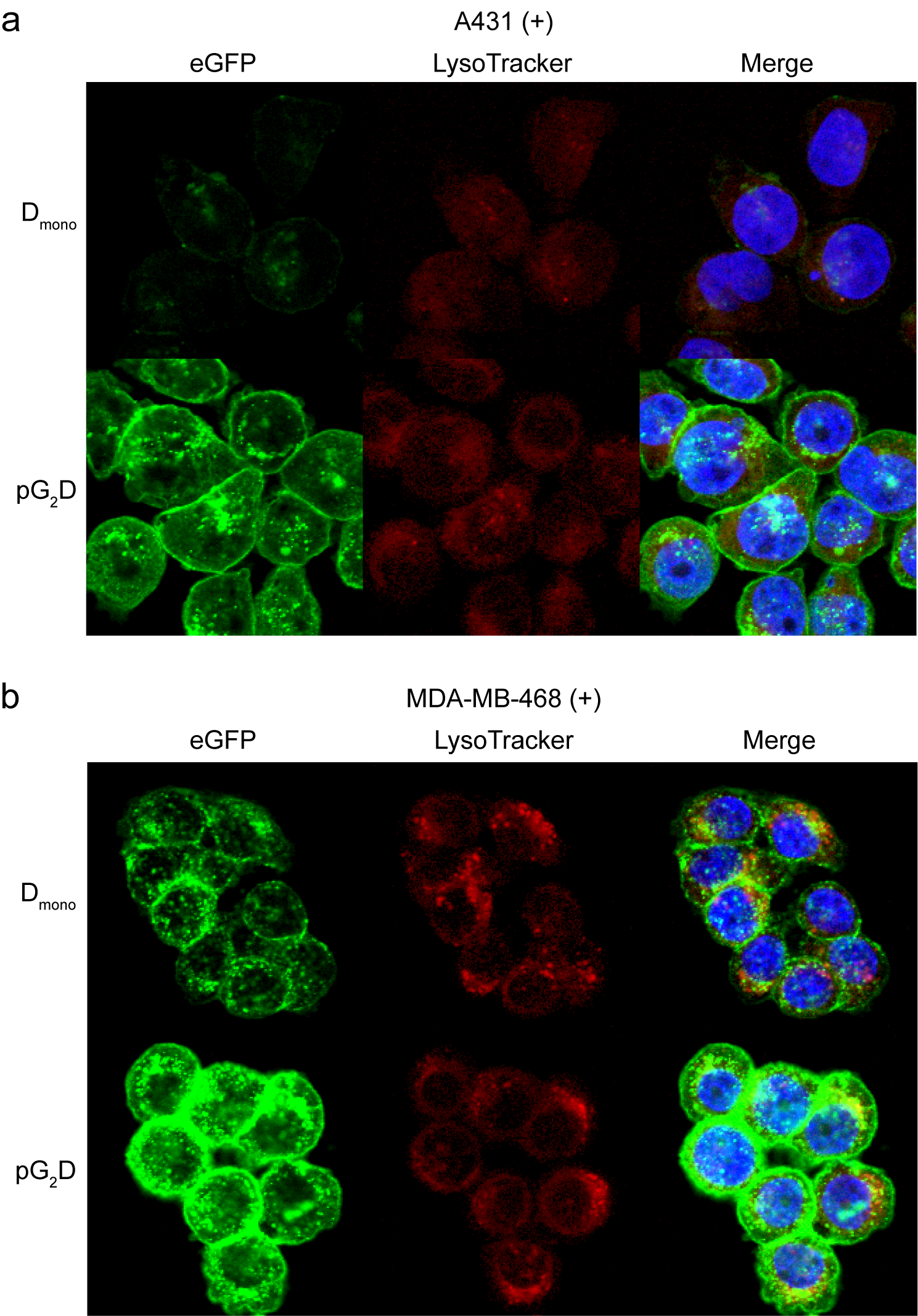

**Figure S10. Enlarged confocal imaging of high EGFR-expression cell lines treated with D_mono_** **and pG_2_D.** **(a)** Confocal image of high EGFR-expressing A431 after treatment with the prodrimers functionalized with the translocation module carrying an EGFR-specific repebody. Intracellular delivery of eGFP was traced with lysotrackers. **(b)** Confocal image of high EGFR-expressing MDA-MB-468 after treatment with the same prodrimers as in (a). All cells were treated for 6 h.

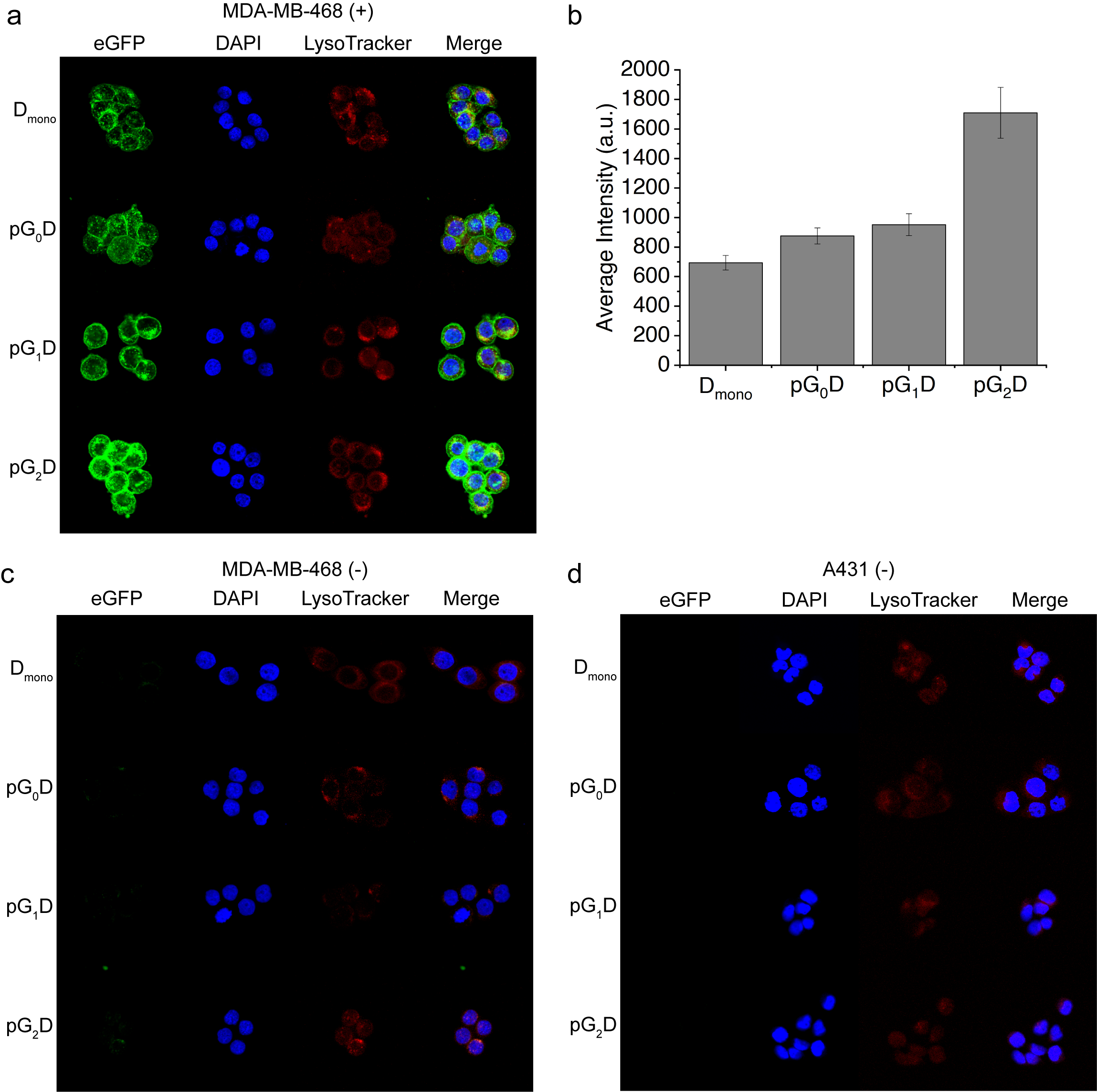

**Figure S11. Confocal images of high EGFR-expressing cell lines treated with the prodrimers functionalized with D_mono_. (a)** Confocal images of high EGFR-expressing MDA-MB-468 cells after treatment with the prodrimers functionalized with the translocation module carrying an EGFR-specific repebody. **(b)** Average cell fluorescence intensity of eGFP from MDA-MB-468 cells treated with the same prodrimers as in (a). **(c)** Confocal images of MDA-MB-468 cells after treatment with the prodrimers functionalized with the translocation module carrying an off-target repebody. (**d**) Confocal images of A431 cells after treatment with the same prodrimers as in (a). All cells were treated for 6 h.

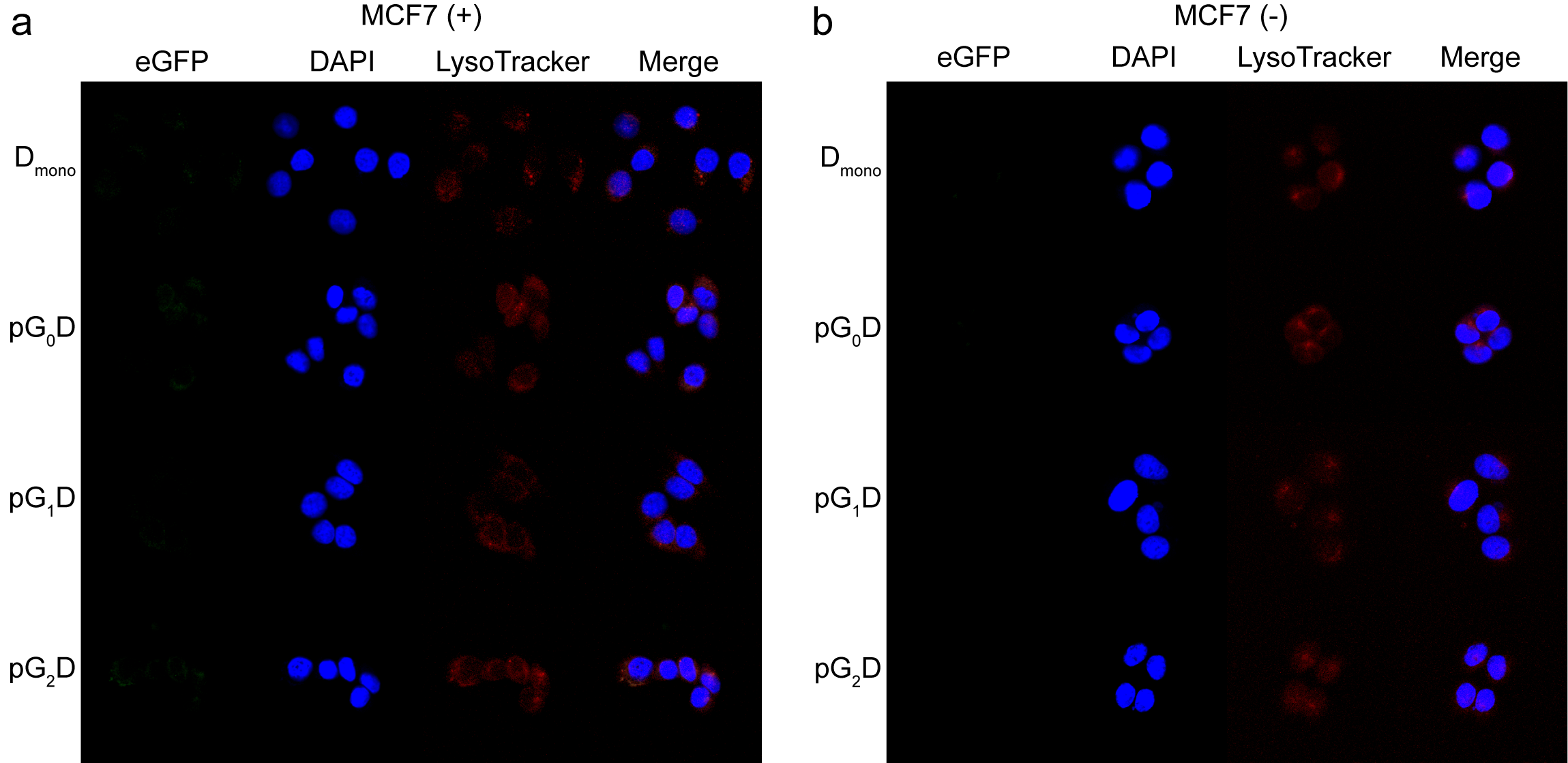

**Figure S12. Confocal images of low EGFR-expressing MCF7 cell line treated with the prodrimers functionalized with D_mono_. (a)** Confocal images of MCF7 cells after treatment with the prodrimers functionalized with the translocation module carrying an EGFR-specific repebody. **(b)** Confocal images of MCF7 cells after treatment with the prodrimers functionalized with the translocation module carrying an off-target repebody. All cells were treated for 6 h.

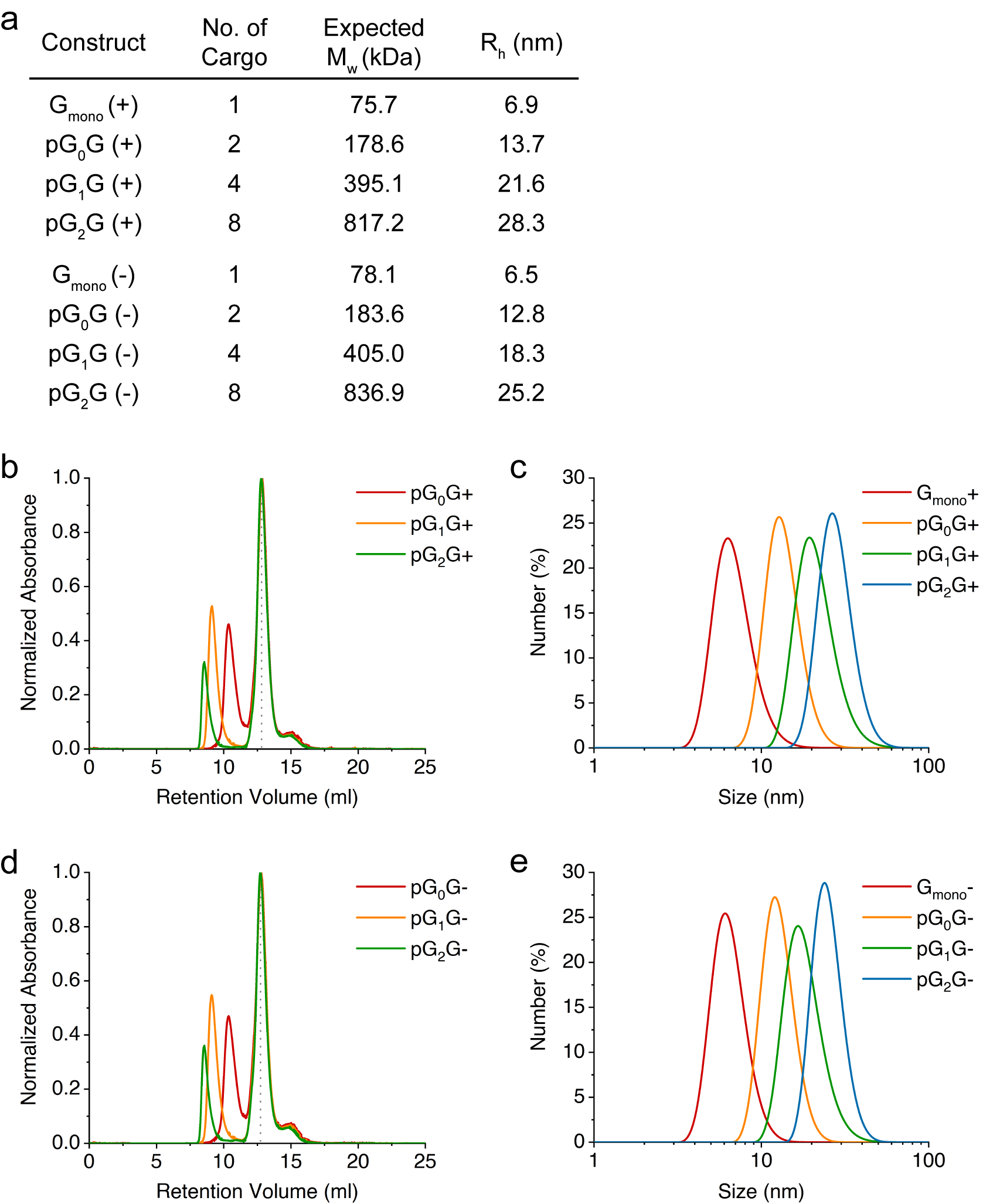

**Figure S13. Biophysical characterization of the prodrimers functionalized with G_mono_. (a)** Summary of the expected molecular masses and the hydrodynamic radii of the prodrimers functionalized with gelonin. (+) and (-) indicate the functionalized prodrimers containing an EGFR-specific repebody and an off-target repebody, respectively. **(b)** Size exclusion chromatography of the functionalized prodrimers containing an EGFR-specific repebody. The peaks represent the normalized absorbance of each prodrimer at 280 nm. The dotted line represent the excess G_mono_ (+) proteins used for functionalization. **(c)** DLS analysis of the functionalized prodrimers containg an off-target repebody. **(d)** Size exclusion chromatography of the functionalized prodrimers containing an EGFR-specific repebody. The dotted line represent the excess G_mono_ (-) proteins used for functionalization. **(e)** DLS analysis of the functionalized prodrimers containing an off-target repebody.

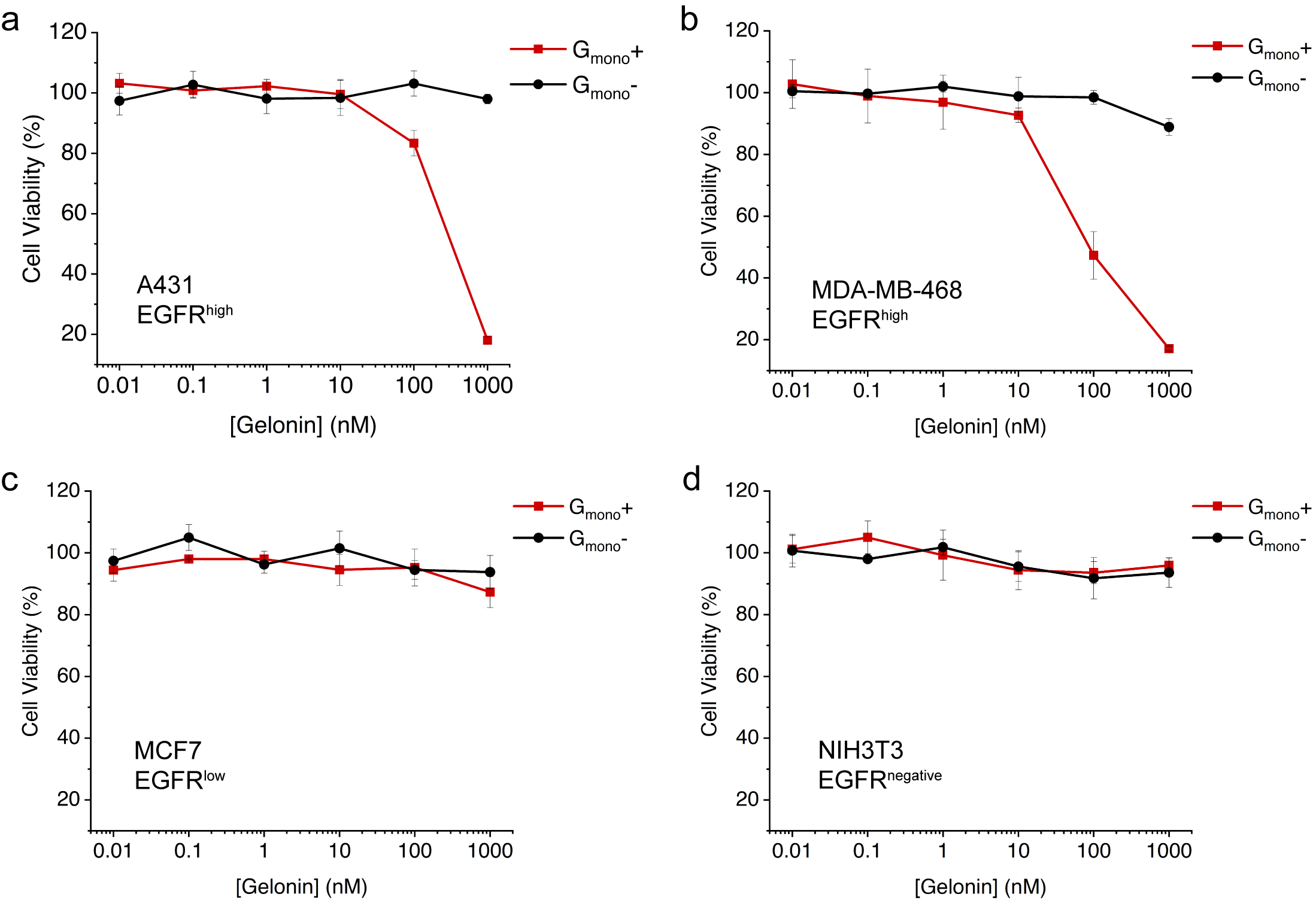

**Figure S14. Viability of cells treated with G_mono_.** Cells were treated with G_mono_ containing an EGFR-specific repebody and an off-target repebody, respectively, for 12 hours, and cell viabilities were measured with respect to the G_mono_ concentration. (+) indicates Gmono carrying an EGFR-specific repebody, while (-) represents G_mono_ carrying an off-target repebody. **(a)** A431, **(b)** MDA-MB-468, **(c)** MCF7, **(d)** NIH3T3.

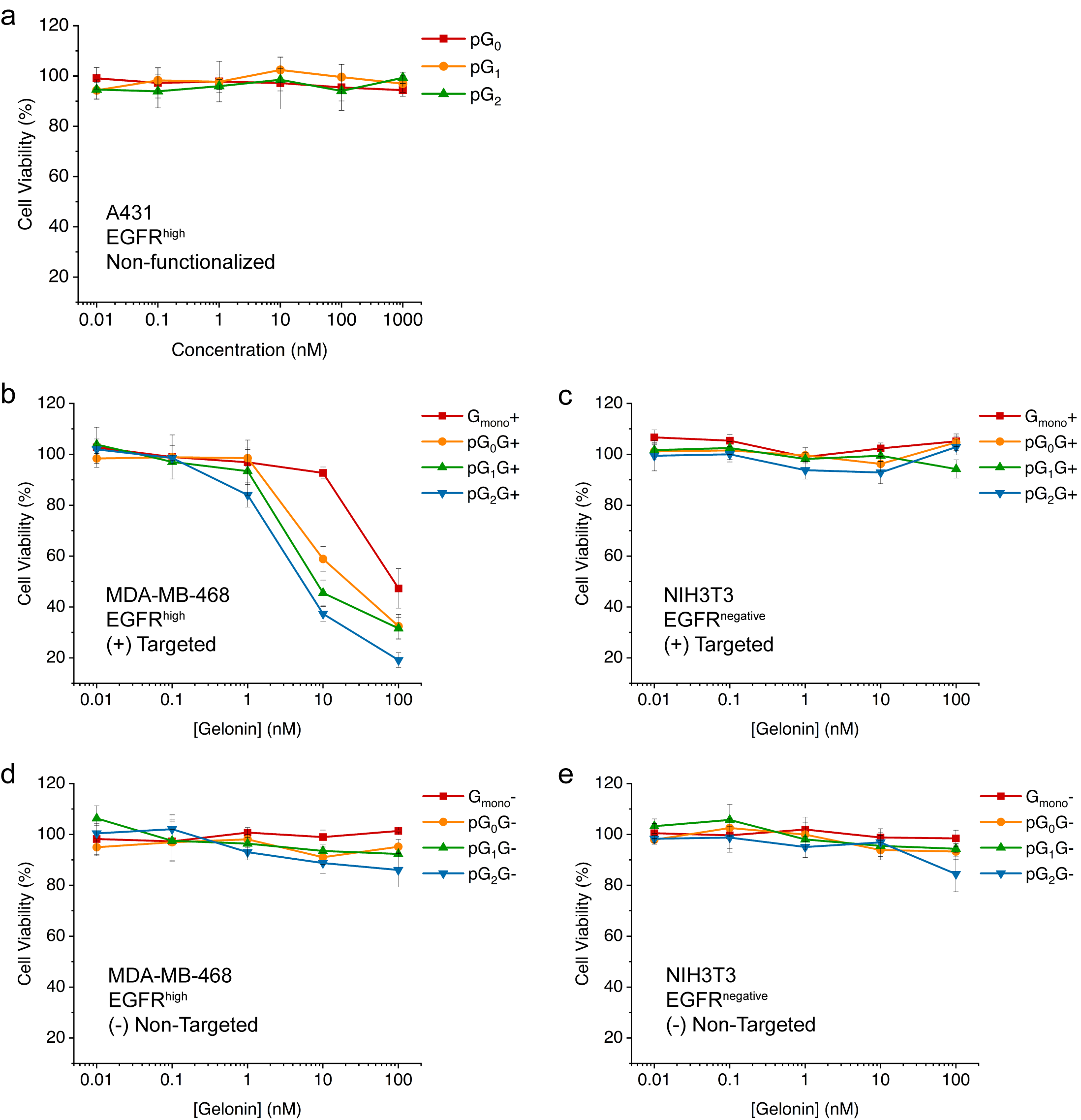

**Figure S15. Viability of cells treated with different generation prodrimers functionalized with G_mono_.** **(a)** Viability of A431 cells treated with different generation prodrimers, pG_0_ to pG_2_. **(b)** Viability of MDA-MB-468 cells after treatment with the prodrimers functionalized with gelonin and an anti-EGFR repebody. **(c)** Viability of MDA-MB-468 cells after treatment with the functionalized prodrimers with gelonin and an off-target repebody. **(d)** Viability of MDA-MB-468 cells after treatment with the functionalized prodrimers with gelonin and an EGFR-specific repebody. **(e)** Viability of MDA-MB-468 cells after treatment with the functionalized prodrimers with gelonin and an off-target repebody. All cells were treated for 12 h.

**Table S1.** Amino acid sequences of the core protein (pG0) and two building blocks (B_1_ and B_2_). Blue and green sequences indicate SnoopCatcher and SpyCatcher, respectively. Blue and green underlined sequences indicate SnoopTag and SpyTag, respectively.

**pG_0_**

MKPLRGAVFSLQKQHPDYPDIYGAIDQNGTYQNVRTGEDGKLTFKNLSDGKYRLFENSEPAGYKPVQNKPIVAFQIVNGEVRDVTSIVPQDIPATYEFTNGKHYITNEPIPPKGSAGSAAGSGEFGGKPLRGAVFSLQKQHPDYPDIYGAIDQNGTYQNVRTGEDGKLTFKNLSDGKYRLFENSEPAGYKPVQNKPIVAFQIVNGEVRDVTSIVPQDIPATYEFTNGKHYITNEPIPPKGGHHHHHH

**B_1_**

MDYDIPTTENLYFQGAMVDTLSGLSSEQGQSGDMTIEEDSATHIKFSKRDEDGKELAGATMELRDSSGKTISTWISDGQVKDFYLYPGKYTFVETAAPDGYEVATAITFTVNEQGQVTVNGKATKGDAHIGSAGSAAGSGEFGGKLGSIEFIKVNKGSAGSAAGSGEFGGDYDIPTTENLYFQGAMVDTLSGLSSEQGQSGDMTIEEDSATHIKFSKRDEDGKELAGATMELRDSSGKTISTWISDGQVKDFYLYPGKYTFVETAAPDGYEVATAITFTVNEQGQVTVNGKATKGDAHIGGHHHHHH

**B_2_**

MKPLRGAVFSLQKQHPDYPDIYGAIDQNGTYQNVRTGEDGKLTFKNLSDGKYRLFENSEPAGYKPVQNKPIVAFQIVNGEVRDVTSIVPQDIPATYEFTNGKHYITNEPIPPKGSAGSAAGSGEFGGAHIVMVDAYKPTKGSAGSAAGSGEFGGKPLRGAVFSLQKQHPDYPDIYGAIDQNGTYQNVRTGEDGKLTFKNLSDGKYRLFENSEPAGYKPVQNKPIVAFQIVNGEVRDVTSIVPQDIPATYEFTNGKHYITNEPIPPKGGHHHHHH

**Table S2.** Amino acid sequences of the constructed proteins used for the functionalization of prodrimer. Blue, green, purple and red sequences indicate an EGFR-specific repebody, eGFP, TDP and gelonin, respectively. Green and blue underlined sequences represent SpyTag and SnoopTag, respectively. For the off-target cargos, the sequence of an EGFR-specific repebody is replaced with that of an off-target repebody.

**T_mono_**

METITVSTPIKQIFPDDAFAETIKANLKKKSVTDAVTQNELNSIDQIIANNSDIKSVQGIQYLPNVRYLALGGNKLHDISALKELTNLTYLMLHYNQLQILPNGVFDKLTNLKELYLSENQLQSLPDGVFDKLTNLTELDLSYNQLQSLPEGVFDKLTQLKDLRLYQNQLKSVPDGVFDRLTSLQYIWLHDNPWDCTCPGIRYLSEWINKHSGVVRNSAGSVAPDSAKCSGSGKPVRSIICPTASGSAGSAAGSGEFGGAHIVMVDAYKPTKGSSKLGSIEFIKVNKGSLENHHHHHH

**F_mono_**

MVSKGEELFTGVVPILVELDGDVNGHKFSVSGEGEGDATYGKLTLKFICTTGKLPVPWPTLVTTLTYGVQCFSRYPDHMKQHDFFKSAMPEGYVQERTIFFKDDGNYKTRAEVKFEGDTLVNRIELKGIDFKEDGNILGHKLEYNYNSHNVYIMADKQKNGIKVNFKIRHNIEDGSVQLADHYQQNTPIGDGPVLLPDNHYLSTQSALSKDPNEKRDHMVLLEFVTAAGITLGMDELYKASGSAGSAAGSGEFGGAHIVMVDAYKPTKGSSKLGSIEFIKVNKGSLENHHHHHH

**D_mono_ (+)**

METITVSTPIKQIFPDDAFAETIKANLKKKSVTDAVTQNELNSIDQIIANNSDIKSVQGIQYLPNVRYLALGGNKLHDISALKELTNLTYLMLHYNQLQILPNGVFDKLTNLKELYLSENQLQSLPDGVFDKLTNLTELDLSYNQLQSLPEGVFDKLTQLKDLRLYQNQLKSVPDGVFDRLTSLQYIWLHDNPWDCTCPGIRYLSEWINKHSGVVRNSAGSVAPDSAKCSGSGKPVRSIICPTEFGGGSGGGSGGGSGGASLAALTAHQACHLPLETFTRHRQPRGWEQLEQCGYPVQRLVALYLAARLSWNQVDQVIRNALASPGSGGDLGEAIREQPEQARLALTLAAAESERFVRQGTGNDEAGAANGSGGGSGGGSGGGSGGTSVSKGEELFTGVVPILVELDGDVNGHKFSVSGEGEGDATYGKLTLKFICTTGKLPVPWPTLVTTLTYGVQCFSRYPDHMKQHDFFKSAMPEGYVQERTIFFKDDGNYKTRAEVKFEGDTLVNRIELKGIDFKEDGNILGHKLEYNYNSHNVYIMADKQKNGIKVNFKIRHNIEDGSVQLADHYQQNTPIGDGPVLLPDNHYLSTQSALSKDPNEKRDHMVLLEFVTAAGITLGMDELYKKDELGGSGSGFLGGGSGSAHIVMVDAYKPTKGSSKLGSIEFIKVNKGSLENHHHHHH

**G_mono_ (+)**

METITVSTPIKQIFPDDAFAETIKANLKKKSVTDAVTQNELNSIDQIIANNSDIKSVQGIQYLPNVRYLALGGNKLHDISALKELTNLTYLMLHYNQLQILPNGVFDKLTNLKELYLSENQLQSLPDGVFDKLTNLTELDLSYNQLQSLPEGVFDKLTQLKDLRLYQNQLKSVPDGVFDRLTSLQYIWLHDNPWDCTCPGIRYLSEWINKHSGVVRNSAGSVAPDSAKCSGSGKPVRSIICPTEFGGGSGGGSGGGSGGASLAALTAHQACHLPLETFTRHRQPRGWEQLEQCGYPVQRLVALYLAARLSWNQVDQVIRNALASPGSGGDLGEAIREQPEQARLALTLAAAESERFVRQGTGNDEAGAANGSGGGSGGGSGGGSGGTSLDTVSFSTKGATYITYVNFLNELRVKLKPEGNSHGIPLLRKKCDDPGKCFVLVALSNDNGQLAEIAIDVTSVYVVGYQVRNRSYFFKDAPDAAYEGLFKNTIKTRLHFGGSYPSLEGEKAYRETTELGIEPLRIGIKKLDENAIDNYKPTEIASSLLVVIQMVSEAARFTFIENQIRNNFQQRIRPANNTISLENKWGKLSFQIRTSGANGMFSEAVELERANGKKYYVTAVDQVKPKIALLKFVDKDPKKDELGGSGSGFLGGGSGSAHIVMVDAYKPTKGSSKLGSIEFIKVNKGSLENHHHHHH

**Off-target repebody (-)**

ETITVSTPIKQIFPDDAFAETIKANLKKKSVTDAVTQNELNSIDQIIANNSDIKSVQGIQYLPNVRYLALGGNKLHDISALKELTNLTYLILTGNQLQSLPNGVFDKLTNLKELVLVENQLQSLPDGVFDKLTNLTYLNLAHNQLQSLPKGVFDKLTNLTELDLSYNQLQSLPEGVFDKLTQLKDLRLYQNQLKSVPDGVFDRLTSLQYIWLHDNPWDCTCPGIRYLSEWINKHSGVVRNSAGSVAPDSAKCSGSGKPVRSIICPT

[1]	X. Chen, J. L. Zaro, W. C. Shen, Adv Drug Deliv Rev 2013, 65, 1357.

[2]	H. Y. Kim, J. A. Kang, J. H. Ryou, G. H. Lee, D. S. Choi, D. E. Lee, H. S. Kim, Acs Chem Biol 2017, 12, 2891.

[3]	J. J. Lee, H. J. Choi, M. Yun, Y. Kang, J. E. Jung, Y. Ryu, T. Y. Kim, Y. J. Cha, H. S. Cho, J. J. Min, C. W. Chung, H. S. Kim, Angew Chem Int Edit 2015, 54, 11020.

[4]	S. C. Lee, K. Park, J. Han, J. J. Lee, H. J. Kim, S. Hong, W. Heu, Y. J. Kim, J. S. Ha, S. G. Lee, H. K. Cheong, Y. H. Jeon, D. Kim, H. S. Kim, Proc Natl Acad Sci U S A 2012, 109, 3299.

[5]	C. J. Crasto, J. A. Feng, Protein Eng 2000, 13, 309.
